# Neural and behavioural manifold dynamics align across interacting individuals

**DOI:** 10.64898/2026.08.19.745562

**Authors:** Atesh Koul, Alessandro Corsini, Francesco Torricelli, Félix Bigand, Sara Abalde, Giacomo Novembre, Alice Tomassini, Alessandro D’Ausilio

**Affiliations:** Dipartimento di Neuroscienze e Riabilitazione, Università degli Studi di Ferrara, Via Fossato di Mortara 17-19, Ferrara, 44121, Italy; Center for Translational Neurophysiology of Speech and Communication, Italian Institute of Technology, Via Fossato di Mortara 17-19, Ferrara, 44121, State, Italy; Neuroscience of Perception and Action, Italian Institute of Technology, Viale Regina Elena 291, Rome, 00161, Italy

## Abstract

Coordinating actions with others is fundamental for social behaviour, requiring the nervous system to continuously adapt motor output to a partner’s evolving behaviour. Yet the neural population principles supporting such coordination remain poorly understood. To address this, we investigated interpersonal coordination across two dual-EEG studies comprising 44 dyads (88 participants) engaged in either instructed finger movement synchronization or spontaneous face-to-face interaction. Combining kinematics-informed deep contrastive learning with dynamical-systems modelling, we identified low-dimensional neural manifolds. These manifolds aligned geometrically and temporally across interacting partners, mirrored their coordinated behaviour, and uncovered interpersonal alignment not captured by traditional synchrony measures. Importantly, these manifolds exhibited flexible attractor-like organization consistent with a synergistic, dynamical account of motor control, with attractor properties that were co-regulated across partners. Collectively, we propose a novel mechanism in which interpersonal coordination emerges through the intermittent updating of internally organized dynamics by a partner’s movements. More broadly, our results establish movement-informed latent-space modeling as a framework for uncovering the low-dimensional population dynamics linking neural activity, neuromuscular control and interpersonal coordination.

## Introduction

Understanding how the nervous system transforms intent into coordinated movement, both within and across individuals, remains a central challenge in neuroscience [1-6]. This problem is compounded by the high dimensionality, redundancy, and variability of the neuromechanical system, whereby multiple neural and mechanical solutions can produce equivalent behavioural outcomes—the so-called degrees-of-freedom problem [7, 8]. Even a seemingly simple action such as reaching to grasp a cup to pass to another individual involves infinitely many combinations of joint angles, muscle activation patterns, and movement trajectories that achieve the same task-level goal [9, 10]. Moreover, motor behaviour is organized across multiple spatiotemporal scales: at a macroscopic level (e.g., hand velocity), movement appears smooth and goal-directed, whereas at a microscopic level, it is decomposed into discrete sub-movements-intermittent velocity pulses potentially reflecting the interaction between neural commands, biomechanics, and sensory feedback [11-16]. Together, neural redundancy, behavioural variability, and multiscale organization raise a fundamental question: how does the nervous system coordinate action despite high redundancy and variability and without explicitly specifying every degree of freedom across spatial and temporal scales?

Despite decades of research, the mechanisms by which this high-dimensional control problem is solved remain poorly understood. Classical accounts posit that the nervous system operates as a predictive controller: it specifies desired movement kinematics, computes motor commands via internal models, anticipates the sensory consequences of action, and minimizes deviations from intended performance [2, 17, 18]. How-ever, such frameworks face fundamental limitations imposed by neural transmission delays, the nonlinear and state-dependent properties of muscles, and the inherent variability of natural movements—particularly in interactive contexts where movements must be continuously coordinated across individuals [19, 20]. Furthermore, predictive frameworks presuppose a given goal against which errors are computed leaving unresolved how interpersonal coordination emerges in less constrained scenarios of spontaneous interaction [21-23]. Taken together, these constraints challenge the extent to which motor control relies exclusively on high-dimensional internal representations and moment-to-moment error-based corrections.

An alternative framework proposes that movement emerges from the synergistic interaction between neural activity, body mechanics, and environmental constraints [24, 25]. Rather than relying primarily on the continuous prediction and correction of detailed movement variables, this view proposes that motor control operates through intrinsic dynamics organized within low-dimensional, synergistic manifolds: control is exerted over task-relevant variables, while biomechanical and reflexive mechanisms contribute to stability and error attenuation. Consistent with this proposal, population-level neural recordings reveal that motor cortical activity evolves along structured, low-dimensional trajectories [26-28]. These low-dimensional trajectories potentially provide a compact basis for generating more complex, time-varying motor outputs without explicitly specifying each degree of freedom [29, 30]. Converging behavioural evidence further indicates that movement patterns at both macroscopic and microscopic scales carry signatures of low-dimensional dynamical structure [31].

Despite growing empirical support, this framework has yet to yield a mechanistic account of how multi-scale movement organization gives rise to coordinated behaviour, particularly during social interactions. Interpersonal coordination poses a fundamental challenge: the partner’s behaviour is variable, continuously evolving and only partially predictable. If it remains unresolved how the nervous system generates predictions about the detailed kinematics of its own actions, it is even less clear how it predicts the behaviour of another agent, who constitutes an independent and dynamically evolving source of variability. Consequently, the mechanisms by which stable interpersonal coordination emerges remain largely unresolved [24, 32].

Here, we test the hypothesis that coordinated action emerges from the dynamic coupling of low-dimensional neural and behavioural manifolds. We further propose that coordination is not continuous but occurs intermittently through movement dis-continuities—submovements— which facilitate information exchange and alignment between interacting systems. Within this framework, interpersonal coordination is achieved not by predicting or prescribing detailed actions, but by bidirectionally shaping and coupling the manifolds that generate them.

We evaluate this framework by jointly analysing neural activity (EEG) and behavioural dynamics (kinematics) during dyadic coordination. Specifically, we demonstrate that: (1) global (whole-brain) interpersonal neural metrics fail to fully capture the neural underpinnings of coordinated behaviour; (2) low-dimensional neural manifolds provide a more accurate characterization of the neural processes that differentiate distinct modes of coordination; (3) dynamics of manifolds better account for both neural and behavioural structure than classical representational models; (4) dynamical manifold properties are correlated across interacting partners; and (5) cross-partner manifold coupling exhibits systematic bidirectional influences, crucially through submovement features. Critically, these organizational principles are observed not only during instructed coordination but also during spontaneous interaction, indicating that they reflect a candidate organising principle observed across two distinct coordination contexts.

By jointly analyzing neural population activity and movement kinematics during coordinated action, we establish a mechanistic link between low-dimensional neural dynamics and multi-scale motor organization. Our findings suggest that coordinated behaviour—spanning fine-grained submovements to macroscopic task-level performance—emerges from the dynamic interaction of manifolds, rather than through the continuous alignment of high-dimensional neural activity.

## Results

### Instructed and spontaneous coordination is evident across macro- and micro-scale kinematics

We characterized interpersonal coordination across macro- and micro-scale movement kinematics in two independent studies. In study 1 (finger synchronization), 21 dyads (42 participants) performed rhythmic index finger flexion-extension movements either in coordination with a partner (Social condition: In-phase and Anti-phase) or alone (Solo condition). Study 2 involved the reanalysis of an independently collected dataset in which 23 dyads (46 participants) engaged in spontaneous, unconstrained face-to-face interaction without explicit movement instructions, contrasted with a No-vision condition in which a visual occluder prevented mutual observation [33].

We confirmed that participants in study 1 exhibited robust coordination across both social conditions (In-phase and Anti-phase) and movement scales (macro and micro). At the macroscopic scale, dyads showed strong positive velocity correlations during In-phase coordination (*M* = 0.76, *SE* = 0.01) and strong negative correlations during Anti-phase coordination (*M* = −0.75, *SE* = 0.01). At the microscopic scale and consistent with previous reports [34, 35], In-phase movements yielded negative submovement velocity correlations (*M* = −0.39, *SE* = 0.02), whereas Anti-phase movements yielded low, positive correlations (*M* = 0.06, *SE* = 0.01). Coordination was significantly stronger during In-phase than Anti-phase movements at both scales (macro: *t*(20) = 73.61, *p* < 0.001; micro: *t*(20) = −14.02, *p* < 0.001), consistent with increased variability and the well-established stability advantage of In-phase coupling [36, 37].

These findings were replicated and extended in study 2. Despite the absence of explicit movement instructions, participants produced weakly but significantly coordinated actions at both macro (*M* = 0.09, *SE* = 0.02) and micro scales (*M* = 0.41, *SE* = 0.05), with interpersonal correlations significantly higher in the Vision compared to the No-vision condition at both scales (macro: *p* = 0.041; micro: *p* = 0.019). Together, these findings confirm that participants coordinated their movements across spatio-temporal scales under both instructed and spontaneous conditions, providing the behavioural foundation for subsequent neural analyses.

### Global interpersonal neural synchrony fails to predict behavioural coordination

We tested whether behavioural coordination could be predicted from high-dimensional neural data using established interpersonal neural synchrony (INS) metrics. For both studies, we computed two canonical INS measures—phase-locking value (PLV) and power envelope amplitude correlation (AmpCorr) —capturing phase-based and linear amplitude coupling, respectively [33, 38]. Metrics were computed for corresponding electrode pairs across dyads and correlated with behavioural synchrony at both macro-scale (velocity) and micro-scale (submovement velocity).

In study 1, cluster-based permutation testing identified a significant PLV cluster differentiating Anti-phase from Solo No-vision condition (*p* = 0.04; see Fig. S1 for results of all contrasts); however, dyad-level neural synchrony changes within this cluster did not correlate with any dyadic behavioural synchrony measure (Spearman correlations, all *ps >* 0.05). In study 2, a significant INS (AmpCorr) cluster emerged for the Vision versus No-vision contrast (*p* < 0.05), replicating prior work [33]. Yet this modulation likewise showed no significant relationship with behavioural coordination at either scale (*ps >* 0.05). This systematic dissociation between global INS and behavioural performance indicates that global synchrony metrics might not fully capture the neural mechanisms directly supporting coordinated action.

### Behaviourally constrained neural embeddings reveal condition-specific interpersonal alignment

To quantify neural alignment driven by behavioural parameters, we projected EEG data into a low-dimensional latent manifold using the contrastive learning frame-work CEBRA [39], with behavioural variables as auxiliary signals (Figure 3A). Two behavioural representations guided the embeddings: (1) macro-scale finger velocity (0.1-20 Hz bandpass), and (2) micro-scale submovement velocity (velocity bandpassed at 2-3 Hz). Embeddings were learned jointly across both partners and all trials within each dyad, yielding a 3-dimensional latent space per trial (*d* = 3).

**Fig. 1.**
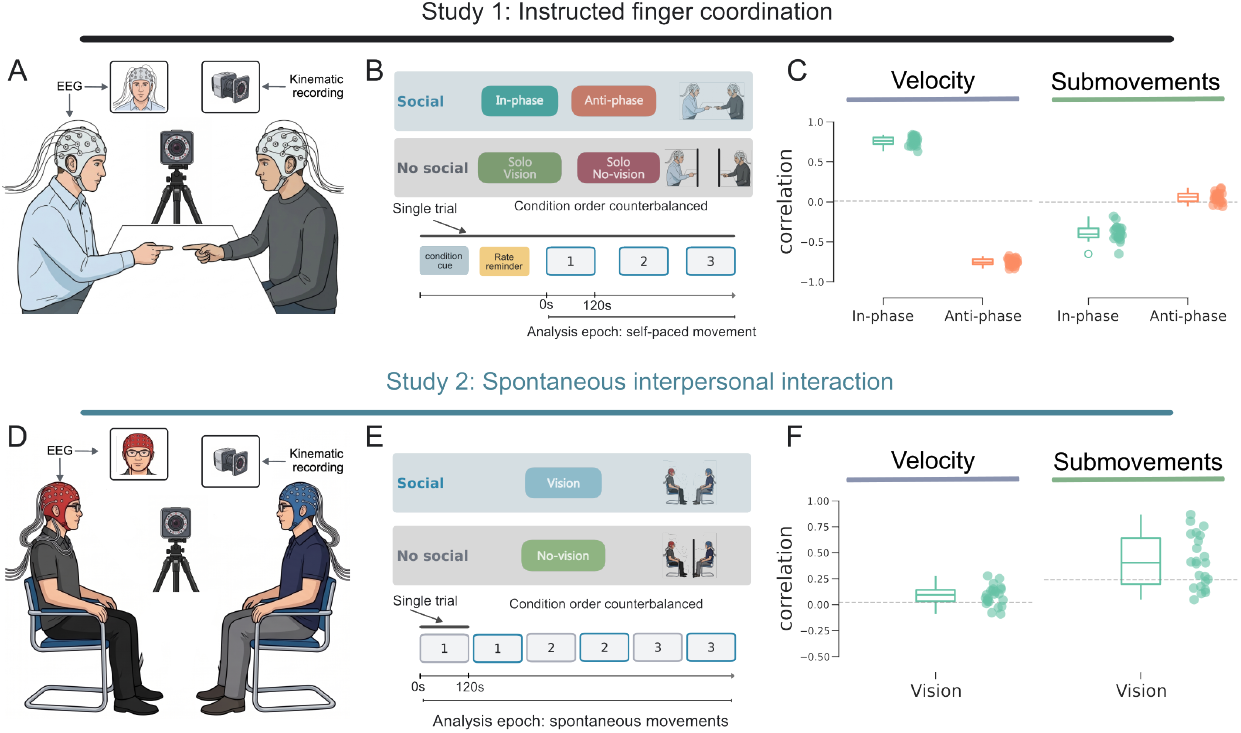
Experimental setup and paradigm. **(A)** Schematic of the dyadic finger synchronization task (study 1). Participants were seated face-to-face across a table while we simultaneously recorded their neural activity (EEG) and tracked their index finger movements (using an infrared motion capture system). The participants were seated at a distance of ≈1m. **(B)** Experimental and trial procedure - Participants were asked to perform rhythmic index finger flexion-extension movements (≈0.25 Hz) either alone (Solo) or in coordination with a partner (Social). In the Social condition, dyads coordinated either In-phase (mirror-symmetric movements) or Anti-phase (same-direction movements). For each condition, we collected three consecutive 2 min trials. Each trial started with instructions on the current condition followed by a finger synchronization rate reminder.**(C)** Coordination results. Participants showed a strong interpersonal coordination both at macro-scale (velocity) as well as at micro-scale (at movement intermittency - submovements). Specifically, at macro-scale, participants showed higher positive correlations during In-phase compared to Anti-phase (*t*(20) = 73.61, *p* < 0.001). At the micro-(submovement) scale, we found a significant negative correlation for In-phase compared to Anti-phase movements (*t*(20) = −14.02, *p* < 0.001). **(D)** Schematic of the spontaneous dyadic interaction task (study 2). Participants sat facing each other and were asked to behave naturally without speaking or making co-verbal gestures. Similar to study 1, neural activity and kinematic data were recorded. **(E)** he experiment comprised two conditions that manipulated interpersonal visual contact: participants could either see one another (Vision) or not (No-vision). For each condition, data were collected across three 2-minute trials, yielding six trials in total. **(F)** Similar to study 1, participants showed greater coordination when they could see each other compared to when they could not. his was so at both velocity (*t*(22) = 2.17, *p* = 0.041) and at submovement scale (*t*(22) = 2.52, *p* = 0.019).

**Fig. 2.**
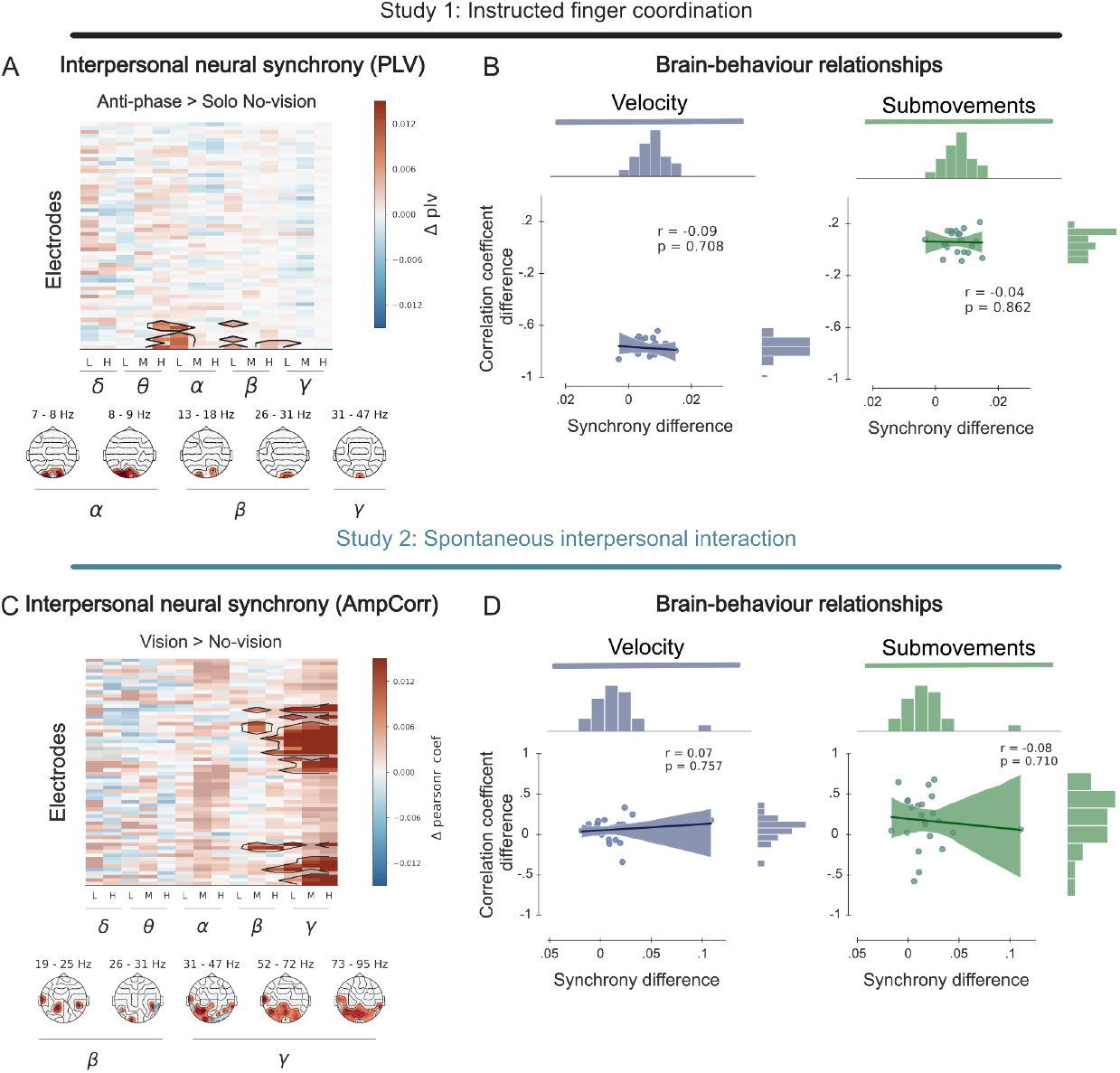
Global interpersonal neural synchrony (INS) does not predict coordination performance. **(A)** opographical maps of significant INS [measured using Phase-locking value (PLV)] in study 1. We find significant INS clusters only for Anti-phase vs. Solo contrasts. **(B)** Scatter plots showing the absence of correlation between neural synchrony (PLV, derived from Anti-phase vs Solo No-vision condition) and behavioural synchrony (macro/micro velocity correlations) across dyads. **(C) T**opographical maps of significant INS (measured using power envelope amplitude correlations) clusters for Vision vs. No-vision contrast in study 2. **(D)** Scatter plots showing the absence of correlation between neural synchrony and behavioural synchrony (macro/micro velocity correlations).

**Fig. 3.**
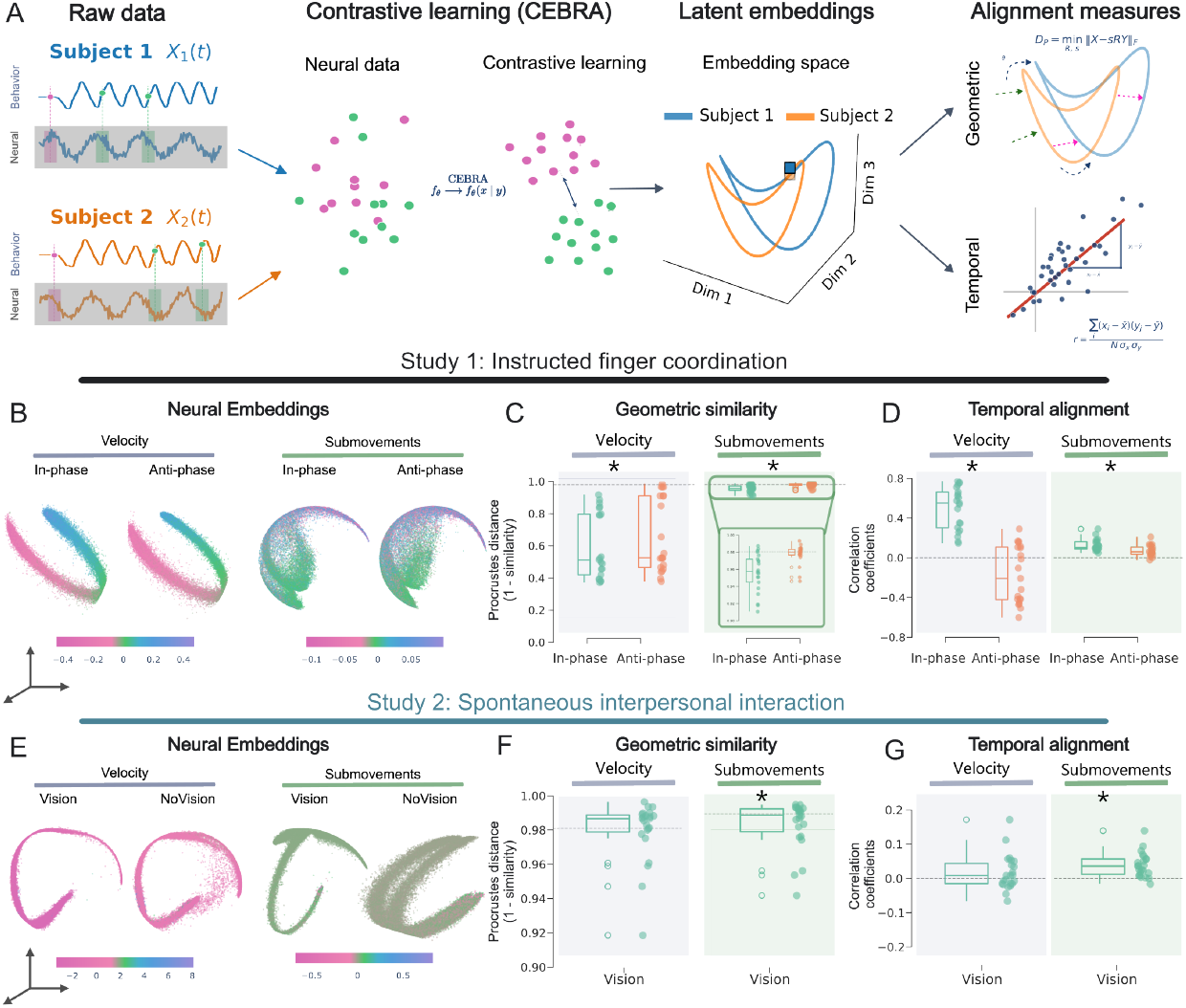
Latent neural embeddings reveal interpersonal alignment. **(A)** Schematic representation of interpersonal alignment computation from low-dimensional neural manifolds. High-dimensional EEG data (**s**_*t*_ ∈ ℝ^64^) are mapped to a low-dimensional latent space (**z**_*t*_ ∈ ℝ^3^) via an encoder *f*_*θ*_. Interpersonal alignment is estimated using geometric (procrustes disparity between interacting partners’ embeddings) and temporal similarity (Pearson’s correlation between the embeddings). **(B)** Example embedding (neural) trajectories using velocity (left) and submovements (right) as auxiliary variables for study 1 (In-phase and Anti-phase movements.) **(C)** Geometric similarity between embedding trajectories is higher (lower procrustes disparity) for In-phase compared to Anti-phase movements at both velocity (*t*(20) = −4.41, *p* < 0.001) and submovement scale (inset; *t*(20) = −5.03, *p* < 0.001), **(D)** Temporal similarity is higher (higher correlations) for In-phase compared to Anti-phase movements at both velocity (*t*(20) = 6.37, *p* < 0.001) and submovement scale (*t*(20) = 4.25, *p* < 0.001). **(E)** Example embedding trajectories using velocity (left) and submovements (right) as auxiliary variables for study 2 (Vision and No-vision conditions). **(F)** Geometric similarity is higher for Vision condition compared to No-vision only at submovement scale (*t*(22) = −2.10, *p* = 0.047) but not at macro scale (*p* = 0.67) (**G**) Temporal similarity is again higher for Vision condition compared to No-vision only at submovement scale (*t*(22) = 2.72, *p* = 0.013, macro scale *p* = 0.57).

Interpersonal alignment in this latent space was quantified via two complementary metrics: (1) Geometric similarity using procrustes disparity, capturing geometric similarity of trajectory shapes independently of amplitude, and (2) Temporal alignment using Pearson’s correlation of embedding time series, capturing temporal alignment (In-phase vs. Anti-phase directionality).

In study 1, latent embeddings differentiated between the modes of coordination and revealed a systematic dissociation between movement direction and scale. For Inphase movements, partners exhibited strong neural alignment in both geometry (low Procrustes disparity) and directionality (positive embedding correlations) at both macro and micro scales. Anti-phase movements showed reduced geometric similarity relative to In-phase at both scales (macro: *p* < 0.001; micro: *p* < 0.001). Temporal alignment for Anti-phase movements showed a negative correlation for macro-scale and a positive correlation for micro-scale.

Robustness was confirmed by multiple control analyses. We did not find condition-specific alignment in either shuffled trial surrogates (*p >* 0.05; see Fig. S2A) or when we trained embeddings on shuffled EEG and behavioural data (*p >* 0.05; Fig. S2B). Condition-specific results also did not generalize to physiological tremor (defined as oscillations in velocity band passed 6-12 Hz; Fig. S2C) and were specific to the task-relevant effector (2-3Hz band passed velocities from the wrist marker did not reproduce the effect; Fig. S2D). Results were also robust to embedding dimensionality as revealed by model fits (*d* = 3 optimal; *d >* 3 yielded equivalent results; Fig. S3). Linear dimensionality reduction (PCA) failed to differentiate In-phase from Anti-phase conditions, confirming that alignment is a property of the behaviourally constrained non-linear embedding rather than a generic feature of EEG topography (Fig. S4A,B).

These findings were replicated in study 2 only for micro scales: embeddings for vision condition exhibited greater geometric similarity (*p* = 0.047) and higher temporal correlations (*p* = 0.013) than No-vision embeddings at micro scales, confirming that latent neural alignment distinguishes coordination conditions. We also replicated the finding that PCA did not differentiate between the Vision and No-vision conditions (Fig. S4C,D).

### Latent neural alignment mirrors the geometry of kinematic signals

To determine the origins of interpersonal coordination in these low-dimensional manifolds and to test whether latent alignment reflects the intrinsic structure of the auxiliary behavioural variables, we applied identical alignment metrics—geometric and temporal alignment—directly to the kinematic signals. Behavioural alignment exhibited clear condition-specificity consistent with the neural embedding results, indicating that contrastive learning built upon coordination-relevant differences already present at the kinematic level. In-phase signals showed high geometric similarity at both macro and micro scales, with positive temporal correlations for velocity and negative temporal correlations for submovements. Anti-phase signals showed reduced geometric similarity at both scales, with negative temporal correlations for velocity and positive temporal correlations for submovements (Fig. S5). This congruence demonstrates that the learned neural manifolds parallel the representational geometry of the behavioural variables used to guide them.

### Macro- and micro-scale neural manifolds encode behaviour via distinct transformations

We next characterized the basis functions linking neural manifolds to kinematic variables. Five candidate transformations were evaluated: Linear (*X*), Absolute (|*X*|), Quadratic (*X*^2^), Cosine (cos *X*), and Sigmoid (*σ*(*X*)). At macro-scale velocity, for both studies we find that non-linear transformations generally better explained the embedding geometry even if the actual transformation was different for study 1 (best cosine) and study 2 (sigmoid) indicating a more non-linear mapping (Fig. 4(B,C)). For micro-scale submovement velocity, we observe consistent results for both studies - absolute value transformations dominated, indicating that the neural manifold encodes sub-movement magnitude, independently of movement direction. This scale-dependent dissociation reveals that the neural representation of submovements is agnostic to movement direction, encoding speed pulses rather than signed velocity. This direction-invariant representation therefore provides a robust measure of movement dynamics across studies, motivating our further focus on submovements in subsequent analyses.

**Fig. 4.**
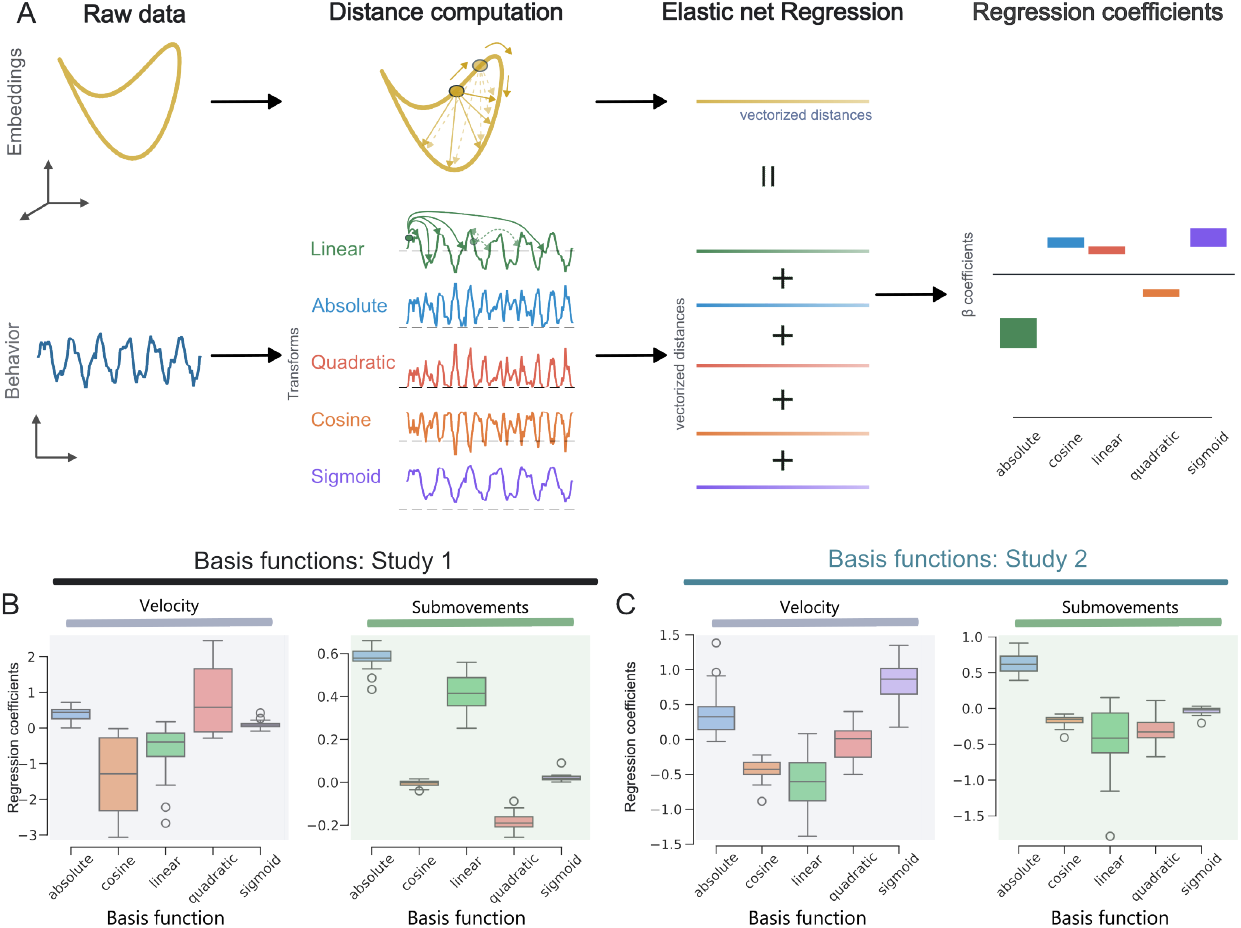
Basis functions linking neural manifolds to kinematic variables. **(A)** Schematic representation of the procedure. Pairwise Euclidean distance matrices were computed for the neural embeddings and a set of five candidate transformations of the kinematic variables, and their upper-triangular elements were vectorized. Standardized transformation distances were used to predict embedding distances using elastic-net regression with 10-fold cross-validation. Coefficient magnitudes were compared using paired t-tests with Benjamini-Hochberg FDR correction. **(B)** Box plots show dyad-averaged, cross-validated elastic-net regression coefficients for Study 1. The cosine transformation produced the largest macroscale coefficient, marginally exceeding the linear transformation (*t*(20) = 1.96, *p* = 0.076) and significantly larger than all other transformations [absolute (*t*(20) = 7.29, *p* < 0.001), quadratic (*t*(20) = 4.66, *p* < 0.001), and sigmoid (*t*(20) = 5.95, *p* < 0.001)]. At the submovement scale, the absolute transformation produced larger coefficients than the linear (*t*(20) = 6.14, *p* < 0.001), cosine (*t*(20) = 42.77, *p* < 0.001), quadratic (*t*(20) = 37.14, *p* < 0.001), and sigmoid (*t*(20) = 44.10, *p* < 0.001) transformations. **(C)** In study 2, the sigmoid transformation produced the largest coefficient at the macroscale, exceeding the linear (*t*(22) = 10.20, *p* < 0.001), absolute (*t*(22) = 3.85, *p* = 0.001), quadratic (*t*(22) = 14.34, *p* < 0.001), and cosine (*t*(22) = 16.44, *p* < 0.001) transformations. At the submovement scale, replicating study 1, the absolute transformation produced larger coefficients than the linear (*t*(22) = 9.30, *p* < 0.001), cosine (*t*(22) = 24.56, *p* < 0.001), quadratic (*t*(22) = 15.13, *p* < 0.001), and sigmoid (*t*(22) = 17.82, *p* < 0.001) transformations.

### Behavioural and neural manifolds exhibit complementary attractor-like dynamics

We next characterized the dynamical structure of behavioural and neural state spaces around submovements (i.e., speed peaks). The behavioural state space was defined as a 2D manifold of position and submovement velocity; the neural state space comprised the *d* = 3 CEBRA embeddings. A family of candidate dynamical models was fit using a rolling autoregressive procedure, providing a unified basis for model comparison by evaluating predictive accuracy of submovement-locked trajectory evolution (Figure 5; Methods). Candidate models comprised: (1) **Trajectory attractor**: attraction toward a time-varying template trajectory **T**(*s*); (2) **Point attractor**: attraction toward a fixed global centroid **c**; (3) **Limit cycle**: oscillatory dynamics in the first two state dimensions; (4) **Error correction**: fractional correction of the initial state toward the global centroid, and (5) **Noise-based control**: a structure-free baseline that predicted the global centroid at each time step.

**Fig. 5.**
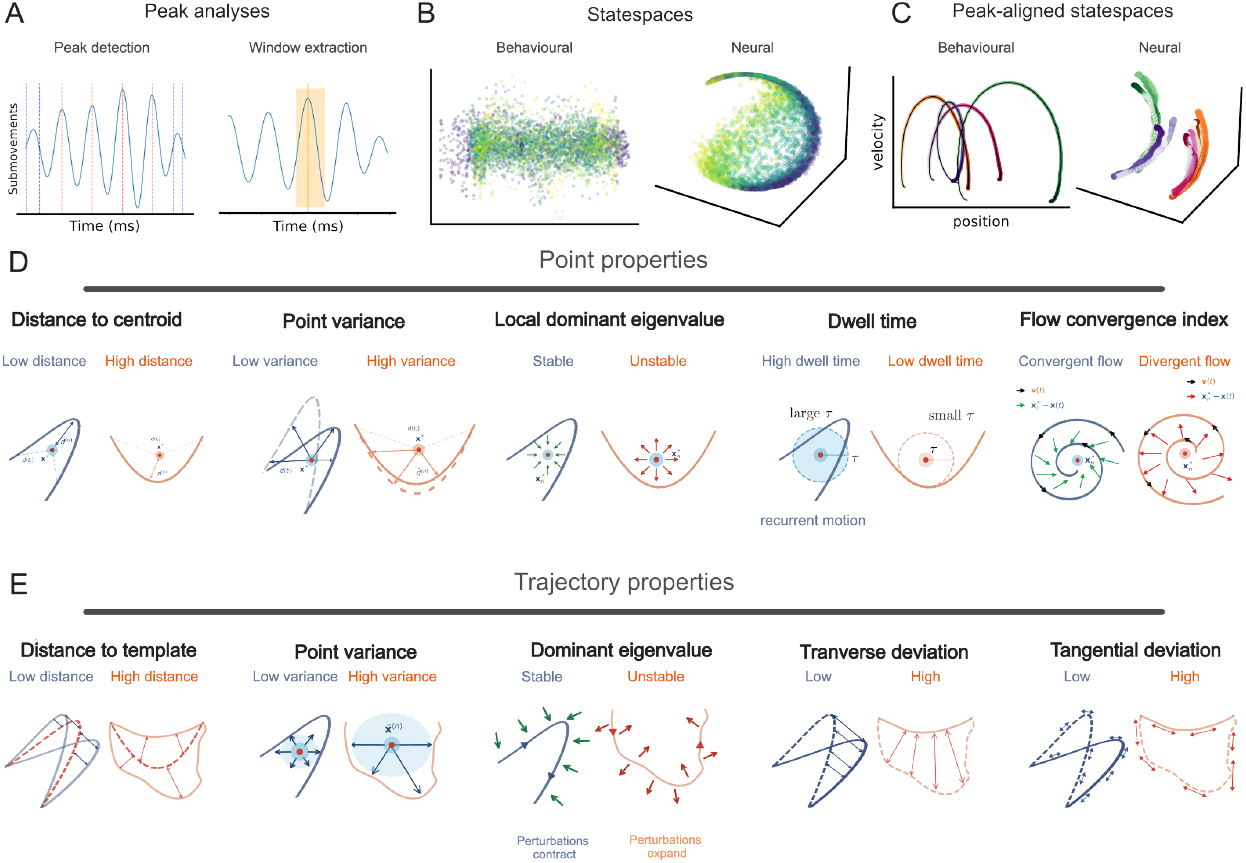
Submovement peak-based analyses. **(A)** Schematic of the submovement peak event based analyses. Submovement peaks were detected via peak detection on velocity profiles (minimum separation 250 ms). Trajectories were resampled to 200 points (*±*200 ms window) for point-to-point geometric comparison. **(B)** Example of the behavioural (position-submovement velocity) and neural (embedding) state spaces. **(C)** Examples of peak-aligned state spaces. **(D)** Schematics of the dynamic point attractor properties of the state spaces. We estimated five properties spanning variability, stability, and convergence. Examples of low and high values are represented below each property. **(E)** Schematics of the trajectory attractor properties. As for point attractors, we defined five properties related to trajectory attractors detailing variability, stability, and convergence.

In study 1 (instructed movements), the trajectory attractor model provided the best account of behavioural dynamics, outperforming point attractor, limit cycle, and control models (Δ*AIC >* 10, Fig. 6A). In study 2 (spontaneous movements), the point attractor model best fit behavioural dynamics (Δ*AIC >* 10, Fig. 6C). In contrast, neural latent dynamics in both studies were best described by point attractor models organized around stable reference states (Δ*AIC >* 10, Fig. 7).

**Fig. 6.**
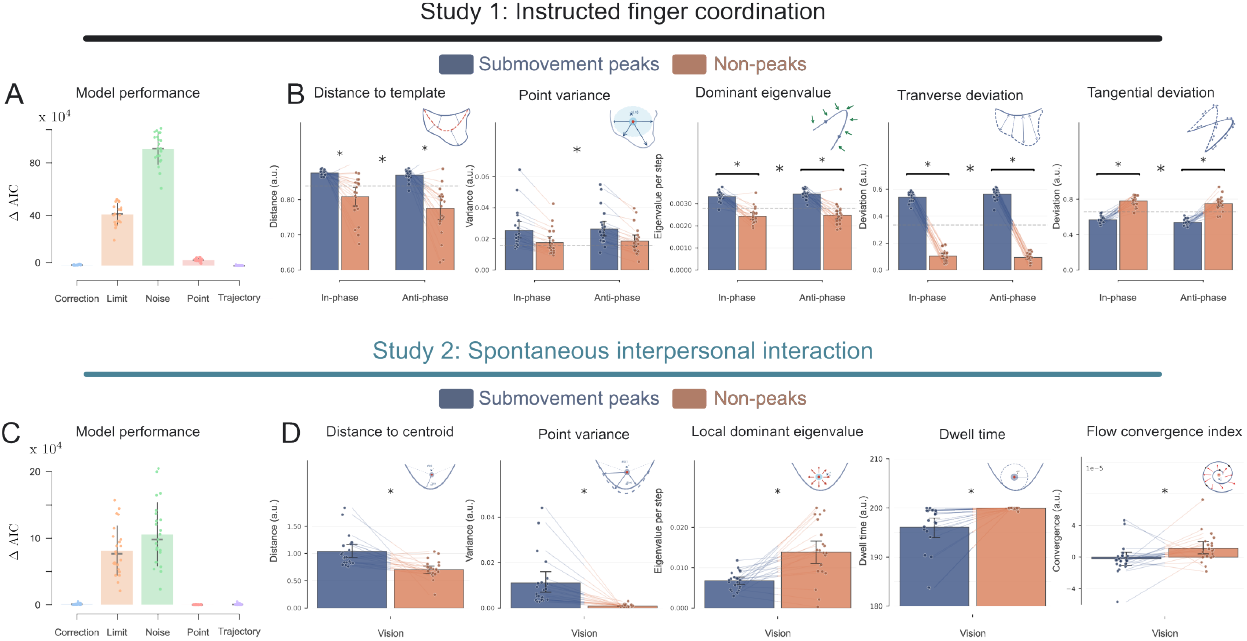
Behavioural state-space properties at submovement peaks vs. non-peaks. **(A)** Bar plots show mean subject-level ΔAIC from rolling autoregressive model comparisons, identifying the trajectory attractor as the best-fitting model (ΔAIC *>* 10 for alternatives). **(B)** Dynamical properties of behavioural state space trajectory attractor for instructed movements. Submovement peaks show weak attractor-like dynamical properties. Specifically, at peaks, we find higher distance to template (Main effect of peak-type *F* (1, 80) = 52.89, *p* < 0.001), higher variance (*F* (1, 80) = 11.33, *p* = 0.001), higher dominant eigenvalue (*F* (1, 80) = 174.76, *p* < 0.001), higher transverse deviation (*F* (1, 80) = 2643.31, *p* < 0.001), and lower tangential deviation (*F* (1, 80) = 314.54, *p* < 0.001). (C) Model comparison for spontaneous movements. Averaged ΔAIC values are plotted for each candidate model. The data were best fit by a point attractor (ΔAIC *>* 10). **(D)** Dynamical properties of behavioural state space point attractor for spontaneous movements. We observe a similar organizational principle (weak attractor) for spontaneous movements as well. Specifically, at peaks, spontaneous behavioural state space demonstrated higher distance to centroid (*t*(21) = 3.99, *p* < 0.001), higher point variance (*t*(21) = 4.16, *p* < 0.001), lower local dominant eigenvalue (*t*(21) = −4.69, *p* < 0.001), a lower dwell time (*t*(21) = −3.66, *p* = 0.002), and a lower flow convergence index (*t*(21) = −2.10, *p* = 0.048). One dyad was excluded in study 2 due to estimation issues. ∗*p* < 0.05, ∗ ∗ *p* < 0.01, ∗ ∗ ∗*p* < 0.001.

**Fig. 7.**
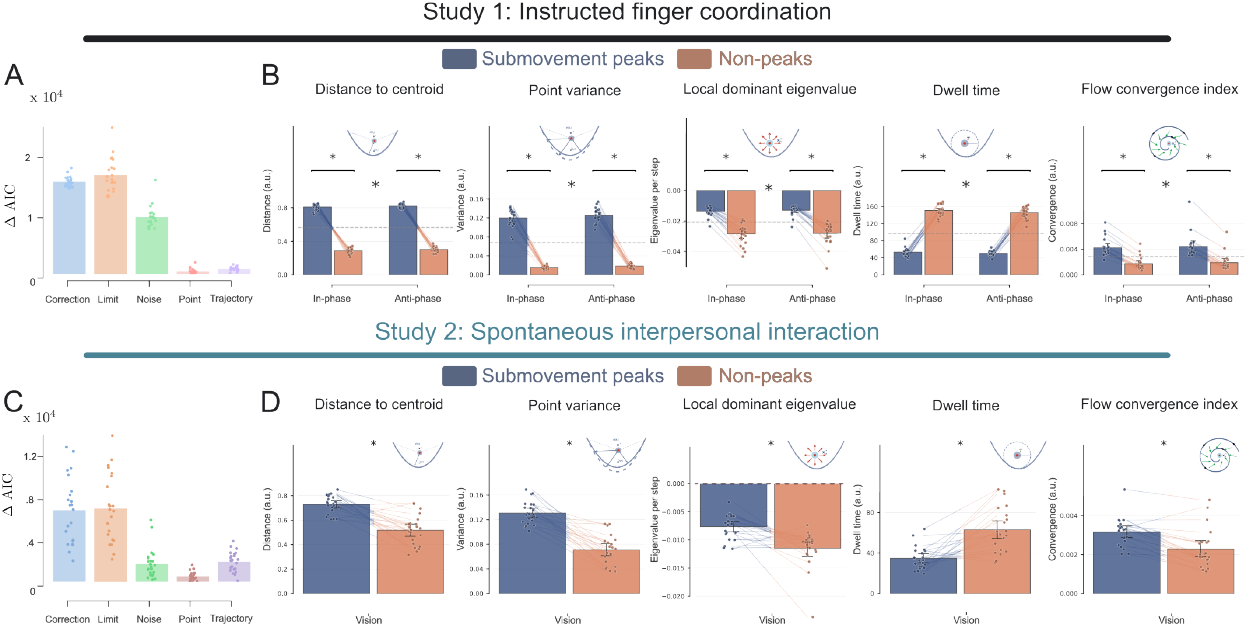
Neural attractor properties at submovement peaks vs. non-peaks. **(A)** Bar plots show mean subject-level ΔAIC, identifying the point attractor as the best-fitting model of the neural manifold (ΔAIC *>* 10 for alternatives). **(B)** Dynamic properties of neural point attractor (study 1). Probing the properties, we again find evidence of a weak attractor parametrized by significantly higher distance to centroid (Main effect of peak-type *F* (1, 80) = 5611.25, *p* < 0.001), higher point variance (*F* (1, 80) = 1570.33, *p* < 0.001), higher local dominant eigenvalue (*F* (1, 80) = 156.89, *p* < 0.001), lower dwell time (*F* (1, 80) = 1671.88, *p* < 0.001), and a higher flow convergence index (*F* (1, 80) = 65.25, *p* < 0.001). **(C)** AIC model comparisons for study 2 reveal a consistent pattern point attractor fits the data better than alternate models (ΔAIC *>* 10). **(D)** Dynamic properties of spontaneous movements fully replicate those of instructed movements with a higher peak distance to centroid (*t*(21) = 6.48, *p* < 0.001), higher point variance (*t*(21) = 9.09, *p* < 0.001), higher local dominant eigenvalue (*t*(21) = 4.38, *p* < 0.001), lower dwell time (*t*(21) = ™5.38, *p* < 0.001), and higher flow convergence index (*t*(21) = −5.96, *p* < 0.001). One dyad excluded in study 2 due to estimation issues. All properties computed for peak-aligned (red) and non-peak (blue) segments. ∗*p* < 0.05, ∗∗*p* < 0.01, ∗ ∗ ∗*p* < 0.001.

Next, we characterized the winning models using a suite of dynamical properties (Tables 1 and 2; Fig. 5(D,E)). At the behavioural level, both studies revealed structured yet flexible organization. During instructed movements, behavioural trajectories diverged more strongly from the mean template at submovement peaks while simultaneously exhibiting reduced stability (higher *λ*^dom^) and increased variability, suggesting that peak states represent transient, organized departures from the underlying trajectory structure (all *ps <* 0.002; Fig. 6B). In study 2, peak states were characterized by greater distance from the attractor reference, increased variability, higher stability, and shorter residence times, consistent with a weakly constrained behavioural regime around submovement events (Fig. 6D).

**Table 1:**
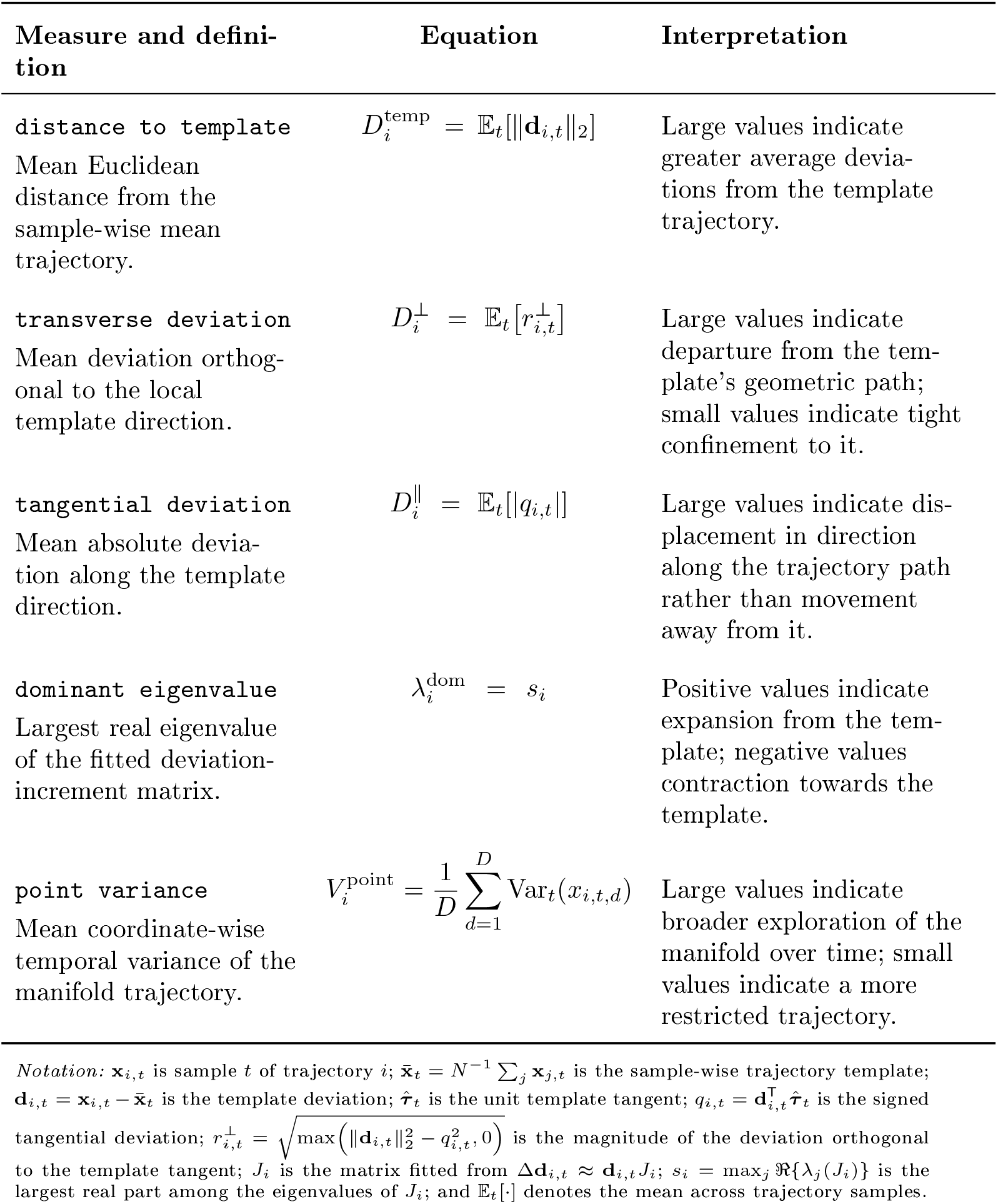
Trajectory-attractor properties.

**Table 2:**
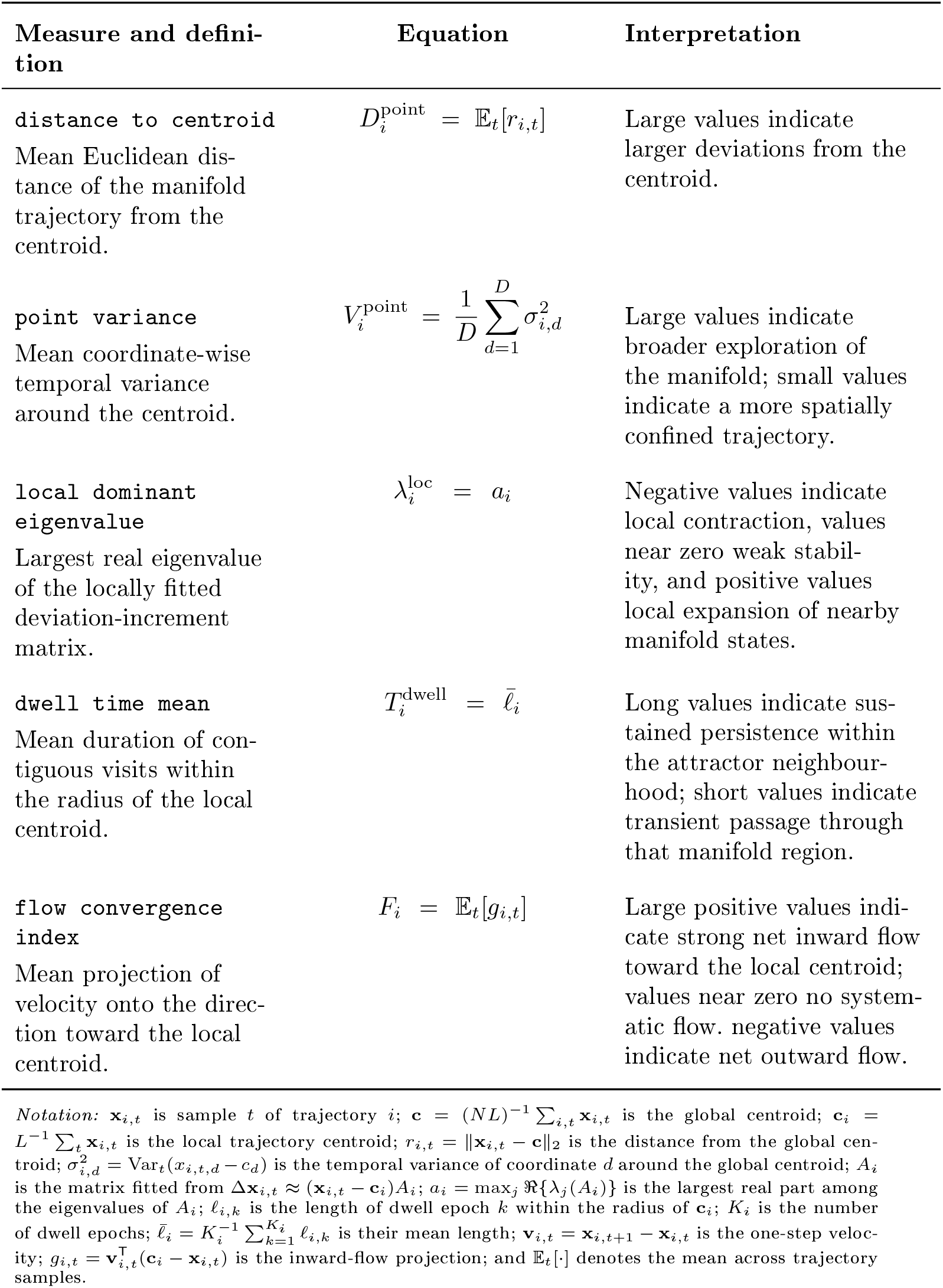
Point-attractor properties.

**Table 3.**
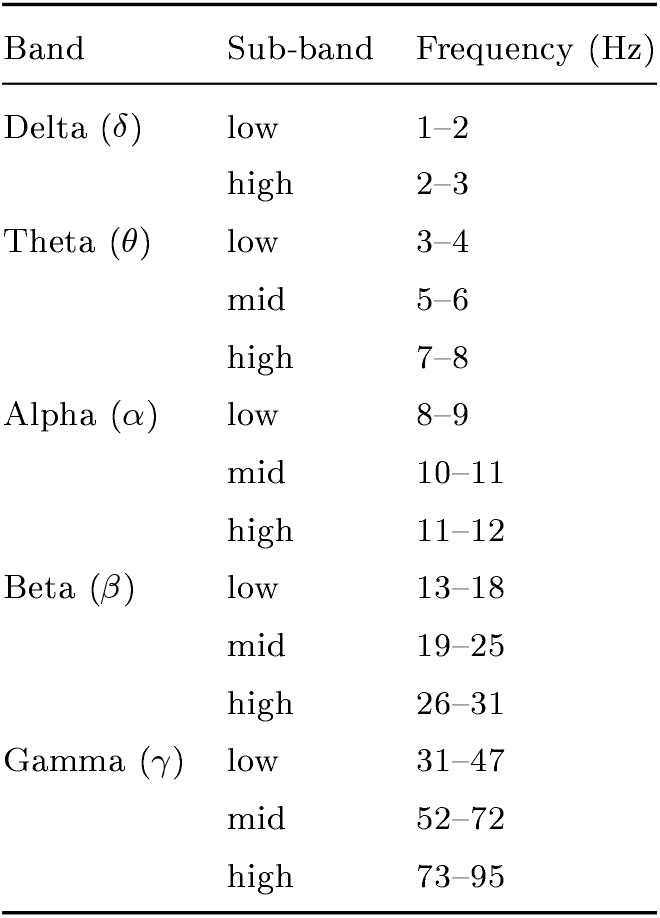
Frequency band and sub-band definitions.

| Band | Sub-band | Frequency (Hz) |
| --- | --- | --- |
| Delta ($\delta$) | low | 1–2 |
|  | high | 2–3 |
| Theta ($\theta$) | low | 3–4 |
|  | mid | 5–6 |
|  | high | 7–8 |
| Alpha ($\alpha$) | low | 8–9 |
|  | mid | 10–11 |
|  | high | 11–12 |
| Beta ($\beta$) | low | 13–18 |
|  | mid | 19–25 |
|  | high | 26–31 |
| Gamma ($\gamma$) | low | 31–47 |
|  | mid | 52–72 |
|  | high | 73–95 |

At the neural level, findings were strikingly consistent across studies: activity moved further from the inferred reference state at peaks while simultaneously exhibiting stronger centripetal flow (higher *F*_*i*_), weaker local contraction (less negative *λ*^loc^), and shorter dwell times (Fig. 7(B,D)). This pattern—transient excursion followed by rapid recovery—is consistent with neural dynamics organized around stable but weakly attracting reference states that support both reliability and flexibility.

Collectively, these results reveal complementary forms of state-space organization: behavioural trajectories become more variable and locally unstable around speed pulses, reflecting adaptive flexibility, whereas neural dynamics transiently depart from and rapidly return to stable reference states. Coordinated interaction may therefore be supported by a balance between behavioural flexibility and neural stability, rather than by rigid deterministic control. In both instructed and spontaneous contexts, behavioural and neural dynamics are characterized by weak, flexible attractor geometries.

### Attractor properties are shared across interacting partners

We next tested whether dynamical properties are correlated across interacting partners. At the behavioural level in study 1, template distance (i.e., sample-wise mean trajectory; 5 and 1) was significantly correlated across partners, with stronger coupling at peaks than non-peaks (In-phase - mean correlation peak = 0.53, non peak = 0.17; Anti-phase - peaks = 0.46, non-peaks = 0.10; Fig. 8). Dynamic time warping (DTW) distance between partners’ trajectories was lower at peaks than non-peaks (In-phase - mean peaks = 0.01, mean non-peaks = 0.16; Anti-phase peaks = 0.037, non-peaks = 0.24), indicating greater behavioural state-space similarity during coordination-critical events.

**Fig. 8.**
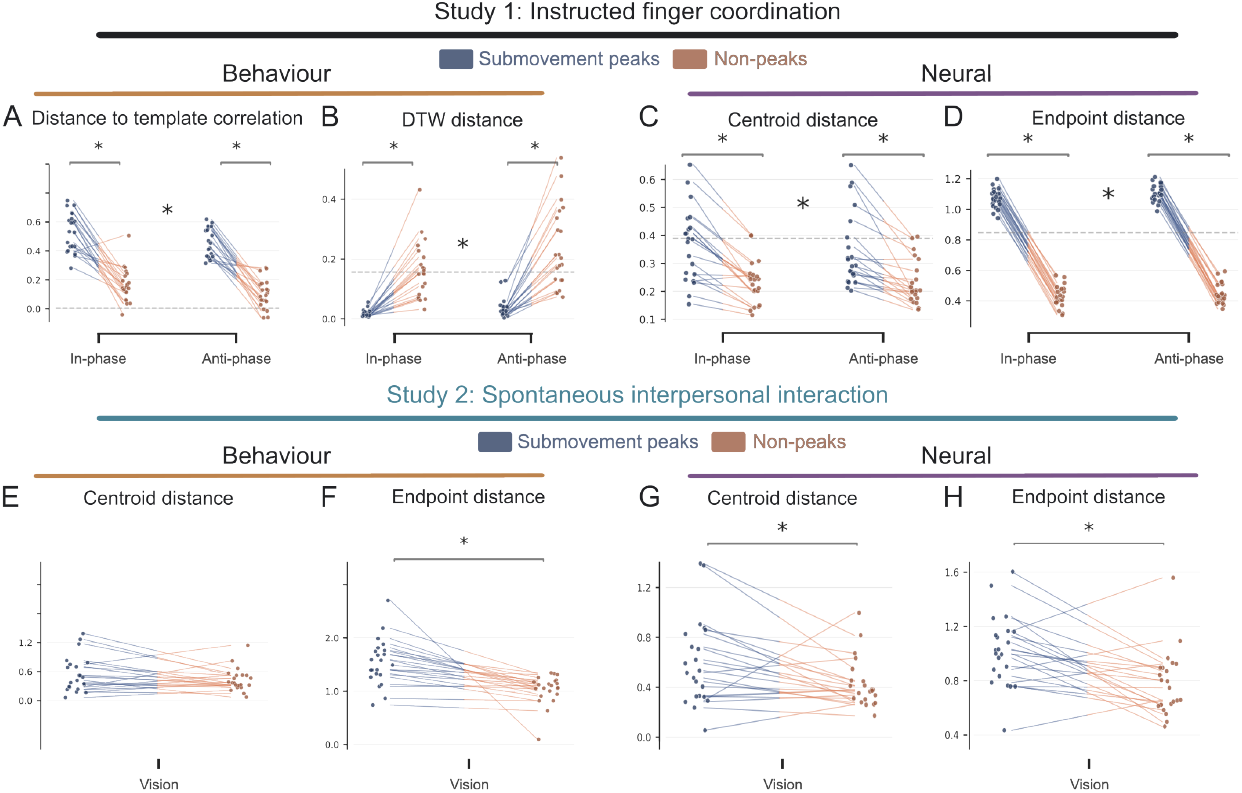
Interpersonal correspondence of attractor properties. **(A)** Study 1: Correlation of template distance between partners (peak vs. non-peak). Paired dot plots representing averaged spearman correlations between distance to template of the two interacting partners. The results show a significantly higher correlation at peaks compared to non-peaks (*F* (1, 80) = 209.33, *p* < 0.001). **(B)** Dynamic time warping distance between partners’ behavioural trajectories (peak vs. non-peak) showed a significantly lower distance at peaks compared to non-peaks (*F* (1, 80) = 87.27, *p* < 0.001). **(C)** Study 1: Paired dot plots representing the neural centroid distance between interacting partners at peaks and non-peaks. Interacting partners showed higher centroid distance during peaks compared to during non-peaks, which showed a higher similarity (*F* (1, 80) = 26.91, *p* < 0.001). **(D)** Study 1: Paired comparison of endpoint distance, defined as the distance between the endpoints of the partners’ neural trajectories, during peak and non-peak segments. Participants that interacted with each other showed a higher distance (lower similarity) between endpoints during peaks compared to non-peaks (*F* (1, 80) = 2096.65, *p* < 0.001).**(E)** Study 2: Centroid distance did not show any significant difference between the peaks and non-peaks (*t*(21) = 1.595, *p* = 0.13) . **(F)** Study 2: End-point distances show a significant difference such that at peaks the end points are more different than non-peaks (*t*(21) = 4.405, *p* < 0.001) **(G)** Study 2: Paired dot plots for neural centroid distance between partners at peaks vs non-peaks. Consistent with the results for instructed movements, we find that centroid distances are higher at peaks compared to non-peaks (*t*(21) = 2.408, *p* = 0.025) **(H)** Study 2: Paired comparison of endpoint distances between partners at peaks vs non-peaks. We find that, consistent with instructed movements, end point distances are higher at peaks (*t*(21) = 3.62, *p* = 0.002) ∗*p* < 0.05, ∗ ∗ *p* < 0.01, ∗ ∗ ∗*p* < 0.001.

In study 2, the pattern reversed: peak segments showed lower interpersonal similarity, evidenced by larger endpoint distances at peaks (Vision - peaks = 1.55, non-peaks = 1.04). This suggests that spontaneous behavioural state space is sustained during non-peak intervals rather than concentrated at transient submovement events.

At the neural level, non-peak intervals showed tighter centroid proximity than peaks (study 1: In-phase - peaks = 0.37, non-peaks = 0.23, Anti-phase - peaks = 0.34, non-peaks = 0.23; study 2: Vision - peaks = 0.60, non-peaks = 0.43). Endpoint distances showed the same pattern (study 1: In-phase - peaks = 1.07, non-peaks = 0.43, Anti-phase - peaks = 1.10, non-peaks = 0.45; study 2: Vision - peaks = 1.02, non-peaks 0.78). This indicates that social interaction induces similar neural attractor geometries across partners, with attractors more tightly clustered during non-peak segments.

Together, these results demonstrate that interpersonal coordination is reflected not only in moment-to-moment behavioural alignment, but also in the correspondence of dynamical invariants—attractor properties—across interacting partners’ brains.

### Behavioural and neural state spaces exhibit bidirectional influences

To quantify directed temporal dependencies between behavioural and neural state spaces, we applied vector autoregressive (VAR) Granger causality analysis [40-42]. For each modality, we defined a joint state vector whose dimensions comprised the state-space coordinates of both partners at each time point. The behavioural joint state space was four-dimensional, comprising position and velocity for each partner, whereas the neural joint state space was six-dimensional, comprising three neural state-space coordinates for each partner. (Fig. 9A). Specifically, the behavioural joint state was defined as 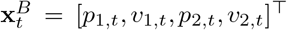, and the neural joint state as 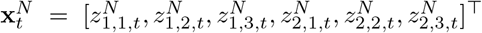. At concatenated submovement peaks, the combined dynamics were best captured by a point attractor geometry, whose properties we assessed next (see also Tables 1 and 2).

**Fig. 9.**
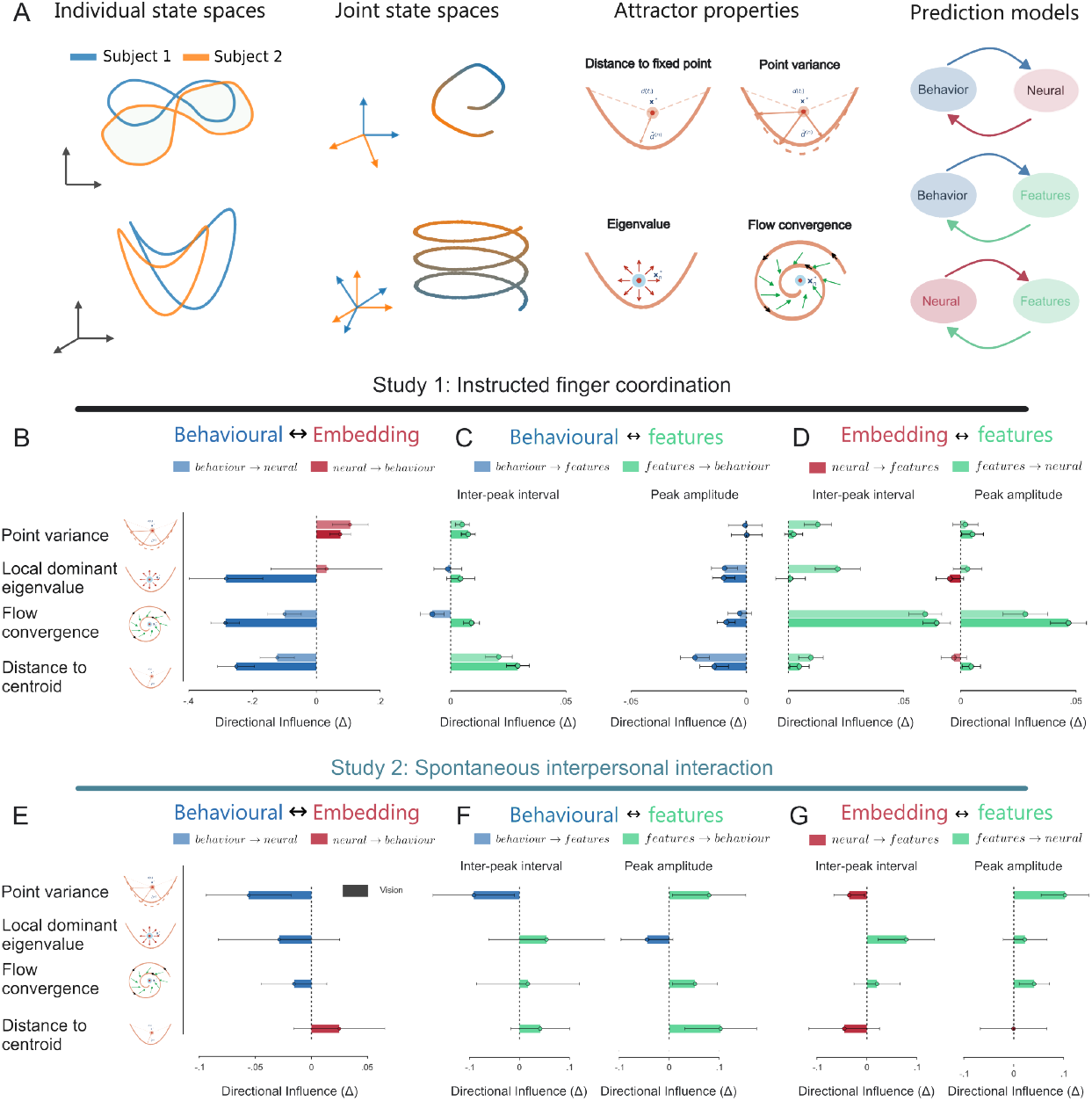
Granger-causal interactions among behavioural state-space geometry, neural dynamics, and submovement parameters. **(A)** Schematic of the Granger causality analysis pipeline. For each dyad, behavioural and neural state-space trajectories were projected in a joint space; model comparison on these combined representations identified a point attractor as the best-fitting geometry (ΔAIC *>* 10). Bidirectional VAR Granger causality tested directed relationships among behavioural attractor dynamics, neural attractor dynamics, and submovement parameters; positive *G*_*X*−*Y*_ indicates stronger *X* → *Y* than *Y* → *X* prediction. (**B**) Directed Granger causality between the behavioural and neural state spaces for point-attractor features. Behavioural attractor dynamics more strongly predicted neural dynamics for distance to the centroid (*F* (1, 80) = 17.85, *p* < 0.001) and flow convergence (*F* (1, 80) = 23.44, *p* < 0.001), whereas the reverse occurred for point variance (*F* (1, 80) = 6.21, *p* = 0.015). **(C)** Directed Granger-causal relationships between behavioural state-space features and submovement parameters in study 1. Inter-peak interval predicted behavioural distance to centroid (*F* (1, 80) = 26.40, *p* < 0.001), whereas behavioural distance, flow convergence, and local dominant eigenvalue predicted peak amplitude (*F* (1, 80) = 11.11, *p* = 0.001; *F* = 4.62, *p* = 0.034; *F* = 4.35, *p* = 0.040, respectively). **(D)** Directed Granger-causal relationships between neural state-space features and submovement parameters in study 1. The inter-peak interval prospectively predicted both ‘distance to centroid’ (*F* (1, 80) = 4.46, *p* = 0.038) and ‘flow convergence’ (*F* (1, 80) = 133.97, *p* < 0.001). Additionally, peak amplitude predicted flow convergence (*F* (1, 80) = 24.19, *p* < 0.001) **(E)** Granger-causality analysis of the behavioural and neural state spaces in study 2. The two state spaces exhibited reciprocal directional influences, with no significant distinction between the two directions. **(F)** Directed Granger-causal relationships between behavioural state-space features and submovement parameters in study 2 revealed mutual influences and no directional effects. **(G)** Directed Granger-causal relationships between neural state-space features and submovement parameters in study 2. Peak amplitude prospectively predicted ‘point variance’ (*t*(21) = → 2.15, *p* = 0.043). One dyad was excluded in study 2 due to estimation issues. ^∗^*p* < 0.05, ^∗∗^*p* < 0.01, and ^∗∗∗^*p* < 0.001.

In Study 1, Granger causality analysis revealed significant directional influences between behavioural and neural state-space dynamics (Fig. 9B). Behavioural distance from the attractor centroid exerted stronger influence on neural state-space organisation than the reverse (*G* _Behav→Neural_ (*D*^point^) *> G* _Behav→Neural_ (*D*^point^)), indicating that moment-to-moment deviations in behavioural attractor geometry prospectively predict neural population dynamics. Neural point variance additionally Granger-caused subsequent behavioural variance (*G*_Neural → Behav_(*V* ^point^) *>G*_Neural → Behav_ (*V* ^point^)), revealing a potentially feedback-like pathway through which neural state variability modulates ongoing joint behavioural organisation. In study 2, both behavioural and neural state spaces mutually influenced each other (Fig. 9E), indicating that spontaneous joint action is characterised by more symmetric mutual coupling, in which behavioural and neural trajectories co-evolve without a dominant direction of influence.

To identify the kinematic-level mechanisms through which these cross-space influences are transmitted, we conducted separate Granger causality analyses linking behavioural and neural state-space properties to discrete submovement parameters. For instructed movements, inter-peak interval significantly predicted behavioural distance from the centroid (Fig. 9C), while behavioural state-space geometry in turn Granger-caused peak amplitude, with behavioural state space flow convergence and local attractor stability further contributing to the prediction of peak amplitude. At the neural level during instructed movements, inter-peak interval prospectively predicted both distance to centroid (Fig. 9D) and flow convergence, demonstrating that submovement timing encodes structural constraints that shape the geometry of neural population dynamics. For spontaneous movements, behavioural state-space geometry and sub-movement features exhibited reciprocal Granger-causal coupling for all features, while at the neural level, peak amplitude prospectively predicted point variance (Fig. 9G), implicating movement amplitude as a key predictor of neural attractor organisation during unconstrained coordination.

Together, these findings demonstrate that interpersonal coupling across attractor dynamics varies with context. In study 1, distinct attractor properties were selectively influenced, while spontaneous coordination in study 2 exhibited reciprocal coupling between behavioural and neural state spaces, with neither direction consistently predominating. Across both studies, discrete submovement parameters provided movement-level pathways linking the two state spaces: inter-peak interval primarily influenced behavioural and neural attractor geometry under instructed conditions, whereas peak amplitude prospectively shaped point variance during spontaneous coordination. These results identify submovement timing and amplitude as context-dependent mechanisms through which behavioural and neural state-space dynamics are prospectively coordinated.

## Discussion

In the present study, we address the fundamental question of how individuals coordinate their actions with those of others. Our findings suggest that interpersonal coordination does not necessarily require a detailed internal representation of either a partner’s kinematics or one’s own. Instead, coordination emerges through the dynamic coupling of low-dimensional neural and behavioural manifolds at fine-grained movement scales. By conditioning latent neural dynamics on behaviourally meaningful (sub)movement events, our approach moves beyond conventional global measures of interpersonal neural synchrony and provides a mechanistic account of how dynamic interplay between behavioural and neural manifolds supports behavioural coordination. These findings have broader theoretical implications for understanding how coordinated action is implemented in the brain without relying on explicit, moment-to-moment representations of high-dimensional movement kinematics.

### Latent neural manifolds go beyond global synchrony metrics

Global interpersonal neural synchrony presumably reflects and aggregates across all simultaneously running neural processes—sensorimotor, attentional, autonomic, task-unrelated—whose coupling with a partner need not be specifically related to movement coordination. This is consistent with the flexible multimodal synchrony framework, which argues that neural synchrony measured globally reflects heterogeneous, parallel processes whose functional relationship to any particular behavioural outcome is diffuse and context-dependent [43, 44].

By conditioning the latent embedding on specific behavioural variables—fingertip velocity and submovement velocity—CEBRA identifies neural dimensions that are maximally informative about the kinematic processes examined [39]. The resulting latent manifolds therefore provide more than a low-dimensional visualisation of neural activity: they retain behaviourally relevant structure that differentiates the coordination mode. In particular, interpersonal alignment within these manifolds was condition-specific and paralleled the corresponding organisation of behavioural coordination. This correspondence suggests that the embeddings recover a task-relevant neural representation of coordination that may be obscured when interpersonal coupling is summarised using global synchrony metrics.

Importantly, the PCA control did not reproduce the condition-specific dissociation observed in the behaviourally conditioned CEBRA manifolds. This finding raises the broader possibility that dimensionality reduction does not necessarily preserve the neural structure relevant to interpersonal coordination, particularly when the reduction is linear and performed without reference to the behaviour of interest (see [45]). Methods such as PCA prioritise dimensions that explain the greatest overall variance in the neural data, but the variance that distinguishes coordination conditions may be comparatively subtle, nonlinear, or distributed across dimensions that account for little global variance. Behaviourally relevant interpersonal structure may therefore be attenuated or discarded during an unconstrained linear reduction, even when it is reliably present in the original neural activity. Our results consequently suggest that the recovery of coordination-related neural geometry depends not only on reducing the dimensionality of the data, but also on how the latent representation is constructed and which information is used to constrain it.

This methodological distinction may help explain why comparable condition-specific neural geometries have not been consistently identified in previous hyperscanning research. Studies relying on global synchrony measures, linear projections, or latent representations derived independently of behaviour may have been insufficiently sensitive to neural structure that is specifically organised around the unfolding kinematics of joint action. Although the present comparison is restricted to PCA and does not establish that all linear or behaviour-independent dimensionality-reduction methods will fail, it motivates the hypothesis that behaviourally constrained and potentially nonlinear embeddings are better suited to revealing task-specific interpersonal neural dynamics. Hyperscanning studies seeking to characterise the neural mechanisms of coordination may therefore benefit from conditioning neural representations on the behavioural processes under investigation, rather than relying exclusively on global coupling measures or unconstrained dimensionality reduction.

### Attractor-like dynamics as a canonical organizational principle

The dynamical framework we find governing both neural latent dynamics and behavioural state spaces is deeply consistent with a broad and growing body of evidence in systems neuroscience. Since Hopfield’s formalization of recurrent network attractors [46], the same general principle—a set of states stabilized through collective positive feedback—has been used to model canonical circuits for motor control, sensory amplification, evidence integration, working memory, decision-making, and spatial navigation [47]. Population recordings in motor cortex have shown that neural activity during reaching evolves along structured, low-dimensional trajectories [26], and that the associated computational motifs are best understood as neural population dynamics rather than as the aggregate of single-neuron tuning properties [48]. The present work extends this framework to the social domain: we show that not only is the neural state space of individual participants organized around point attractor dynamics, but that these attractors become geometrically aligned across interacting brains as a function of social interaction.

A critical property of the observed attractors is their weakness. Local dominant eigenvalues closer to zero, brief dwell times, and strong centripetal flow at sub-movement peaks characterize a system that permits transient local expansion but subsequently undergoes strong centripetal recovery. The system therefore returns reliably to its reference state while remaining readily susceptible to perturbation by incoming behavioural signals. Such properties potentially allow for flexibility especially during mutual adaptation when the behaviour of the partner is not fully predictable. If neural attractors were strongly constraining, the system would resist the perturbations imposed by a partner’s evolving movement dynamics. Instead, if they were absent, coordination would depend on explicit kinematic specification and continuous error correction, which might not allow for a smooth coordinated interactive behaviour. This leads to a testable hypothesis: impaired coordination, including in clinical populations, is associated with more rigid attractor dynamics that constrain flexible adaptation to a partner’s evolving behaviour.

Finally, this view resonates with the equilibrium-point hypothesis of Feldman and Latash [24, 49], in which the controlled variable is not a detailed muscle activation pattern but a threshold configuration of the neuromuscular system—an equilibrium from which movement emerges through the elastic and reflexive properties of the body. The attractor states identified here could potentially reflect a similar process at the neural level. In this view, the neural activity would evolve on a lower-dimensional geometry and observation of partner submovements would reflect potential forms of transient, structured departures. This is consistent with the idea that our nervous system might not encode precise muscle commands but abstract patterns that can be updated flexibly rather than being a rigid trajectory representation [7, 8].

### Submovements as intermittent coupling events

Probing into the mechanisms of neural coordination, we find that submovement peaks do not follow a kinematic noise-like profile but could potentially reflect salient events during interpersonal coordination. This proposition is supported by evidence that movement intermittency is a general phenomenon, documented across human reaching and tracking, non-human primate wrist movements, head movements, and isometric tasks in both humans and non-human primates [11, 35, 50]. This cross-species prevalence indicates that submovements might reflect a fundamental organizational principle of the neuromechanical system. We propose that these events, while they might not have evolved primarily for social signaling, could act as transient opportunities for interpersonal coupling. At submovement peaks, neural activity transiently departs from its attractor reference, centripetal flow is strongest, and—in study 1—behavioural state-space similarity between partners is highest. Peaks thus potentially constitute windows of amplified bidirectional influence: the partner’s sub-movement could be a visible, salient perturbation to an otherwise stable system, and the rapid neural recovery that follows constitutes a form of alignment to the perturbation source.

Our results draw attention to the function of movement intermittency. While these intermittent events have been primarily thought to reflect limitations of the muscle effectors, or of a corrective nature, growing evidence suggests that they might play an important role during interpersonal coordination as well [34, 35]. Our results extend these findings in two principal ways. First, they identify properties of latent spaces that might facilitate coordination - they are agnostic to direction and are encoded through an absolute transform. This suggests that what might be shared during interpersonal coordination is potentially kinematic intensity, not directional specification. Partners potentially share the amplitude and timing of submovement events; the macroscopic trajectory integrates these signals into coherent phase-specific coordination. Second, and more directly, results from temporal dependencies between behavioural-neural state spaces reveal bidirectional influences through peak-amplitude and inter-peak intervals.

More generally, our results are consistent with accounts of a recurrent, bidirectional brain-body system [25, 49]. They suggest that intermittent submovement parameters may provide a mechanism through which neural and bodily dynamics integrate information about a partner’s behaviour and reciprocally update ongoing coordination.

### Compatibility with predictive accounts of motor control

The current low-dimensional dynamical framework is not incompatible with predictive or corrective motor control frameworks. Internal models, forward predictions, and error-correction mechanisms are likely to be essential during unpractised movements, novel skill acquisition, when dynamical policies may not yet have stabilized, and movements performed in unpredictable environments, where sensorimotor delays make purely reactive control insufficient [2, 17]. For well-learned, everyday movements of the kind studied here, however, we suggest that participants rely on motor strategies or policies characterized by flexible yet stable dynamical landscapes. Importantly, this view does not imply a fixed set of geometries, nor does it preclude alternative landscapes that may be adopted in response to perturbations. Predictive processes could instead operate at the level of attractor geometry, shifting the reference configuration in anticipation of an upcoming movement phase rather than recomputing detailed effector trajectories at each time step.

Under this hierarchical account, a partner’s submovement could act as a perturbation to the attractor reference state, modifying the neural geometry without requiring explicit prediction of the partner’s future kinematics. Corrective mechanisms would remain present. Indeed, although our error-correction model was systematically out-performed by the attractor models, the difference was modest. Such mechanisms may operate at lower levels of the sensorimotor hierarchy, correcting moment-to-moment deviations within the envelope defined by attractor dynamics. This interpretation is consistent with accounts that reconcile equilibrium-point control with internal-model frameworks by assigning them to different levels of a control hierarchy [24]: high-level controllers set equilibrium parameters, whereas lower-level spinal and proprioceptive mechanisms correct deviations within those parameters. If correct, this framework generates several testable predictions. For instance, during novel motor-coordination tasks, in which stable dynamical policies have not yet been established, corrective models should outperform dynamical geometric models.

### Conclusion

How does the nervous system coordinate action without explicitly specifying every degree of freedom across spatial and temporal scales? Our findings suggest that such explicit specification may not be necessary. Instead, coordination emerges when the low-dimensional dynamical manifolds that generate movement operate within a regime of weak and flexible attractors. The geometry of these attractors is shaped by the social context and intermittently perturbed at submovement events, creating brief windows during which the state of each system could be influenced by that of the partner. Conventional global synchrony measures fail to capture this mechanism because they aggregate coupling across heterogeneous neural processes, obscuring the specific geometry of movement-relevant neural manifolds. Once this geometry is recovered, interpersonal coordination emerges as a property of coupled dynamical systems, supported by sensorimotor population dynamics that flexibly align with and respond to a partner’s variable and evolving state.

## Methods

### Participants

Data were pooled from two independent studies comprising 88 participants in total. Study 1 (finger synchronization): 42 participants (26 female; mean age 24.1 years, range 20-35) formed 21 dyads. Study 2 (spontaneous interaction): 46 participants (26 female; mean age 21.4 years, range 18-30) formed 23 dyads. Data from study 2 have been partially reported previously [33]. All participants had normal or corrected-to-normal vision and no history of neurological or psychiatric disorders. Written informed consent was obtained from all participants prior to testing, and monetary compensation (25 euros) was provided. Experimental protocols were approved by the respective local ethics committees and conducted in accordance with the Declaration of Helsinki (World Medical Association, 2008). Sample sizes were determined a priori based on prior interpersonal neural synchrony studies [38, 51, 52] rather than formal power analysis.

### Experimental design and procedure

Both experiments employed a within-dyad design manipulating the presence of social interaction during simultaneous dual-EEG and motion capture recordings (Fig. 1A). The Social condition allowed mutual visual access to the partner’s kinematics (finger movements in study 1; whole body in study 2). The control condition (Solo in study 1, No-vision in study 2) prevented visual contact via an opaque occluder placed between participants.

#### Study 1 (Finger Synchronization)

In the Social condition, participants sat facing each other across a table (1 m apart). An occluder panel concealed faces while preserving finger visibility. In Solo conditions, participant’s vision of their own finger was either prevented (Solo No-vision) or allowed (Solo Vision). The task required rhythmic index finger flexion-extension at ∼0.25 Hz. In the Social condition, two coordination modes were used: In-phase (mirror-symmetric: one participant flexes while the other extends) and Anti-phase (same-direction: both participants flex or extend simultaneously). Given the mirror-symmetric seating and hand posture, In-phase required opposite movement directions whereas Anti-phase required same-direction movements. Participants practiced with a metronome (∼1 min) prior to each condition; the metronome was silenced during data collection (Fig. 1B). Each condition comprised 3 trials of 2 min duration (12 trials total per dyad). Condition order was counterbalanced across dyads.

#### Study 2 (Spontaneous Interaction)

During the Social condition, participants sat facing each other at a distance of 3 m and were asked to simply relax and act spontaneously while looking at each other without verbal communication or co-verbal gestures. Participants were not required to necessarily look at each other’s faces or eyes, but rather they were generally asked to look at the body of their partner. In the No-vision condition, an occluder prevented mutual observation. In the original study, participants also completed another condition with a spatial proximity of 1 m. Here, we only analysed the conditions associated with 3 m spatial proximity because those associated with 1 m proximity yielded noisy Vicon estimates (presumably because when the two bodies were too proximal to each other, the visibility of the physical markers was occasionally obstructed). Each condition comprised 3 trials of 2 min (6 trials total per dyad; Fig. 1E). The total session duration was approximately 2.5 hours including setup, calibration, and debriefing.

### Data acquisition and synchronization

Neural activity (dual-EEG) and body kinematics (motion capture) were recorded simultaneously from both participants. System synchronization was achieved via hardware triggers.

**Study 1:** Synchronization used TTL pulses transmitted via parallel port to all acquisition systems, controlled via the Psychophysics Toolbox [53] in MATLAB.

**Study 2:** Synchronization employed the Multi-device Inter-Synchronizer (MIS; https://github.com/ateshkoul/Multi-device-Inter-Synchronizer), a Python-based system (PySimpleGUI) that manages participant metadata (Ethernet), trigger distribution (National Instruments DAQ and virtual serial port), and device coordination across the dual-EEG and motion capture systems.

#### Kinematic recordings

**Study 1:** We recorded three-dimensional marker trajectories (mediolateral *X*, anteroposterior *Y*, vertical *Z*) using a 10-camera Vicon system (300 Hz). Participants wore light-weight retro-reflective hemispheric markers (participant 1: 3 markers, participant 2: 4 markers; see Fig. 1A). The markers were placed on the right hand: distal phalanx of the index finger, metacarpophalangeal (MCP) joint, radial styloid process, and elbow (only for participant 2). For the present analyses, we used the mediolateral (*X*-axis) trajectory of the index-finger marker (as this was the axis where the participants were required to coordinate).

**Study 2:** An 8-camera Vicon system (250 Hz) captured full-body kinematics. Participant 1 wore 18 markers; Participant 2 wore 19 markers across multiple body parts - head, torso, bilateral shoulders, elbows, wrists, knees, feet ([23, 33]). For comparability with study 1, in the present analyses, only the right wrist marker was used.

#### Neural recordings

**Study 1:** We used a 64-channel EEG (Brain Products actiCAP; EasyCap M1 layout; 1000 Hz) system to record the neural activity. Four additional electrodes (FT9, FT10, PO9, PO10) recorded horizontal and vertical electrooculograms (EOG).

**Study 2:** A 64-channel EEG (Ag/AgCl active electrodes; BioSemi ActiveTwo; 2048 Hz) system was used to capture neural activity. The electrodes were placed according to the extended international 10-10 system. One additional electrode at the right outer canthus recorded EOG.

### Data analysis

#### Kinematics preprocessing

**Study 1:** For each trial, marker trajectories were visually inspected for labeling errors and data gaps. Gaps were interpolated in Vicon Nexus using smooth quintic spline interpolation (because of the regular nature of the task). Position was taken as the x-dimension and velocity was computed as the first time derivative.

**Study 2:** Gap filling was performed using Pattern Fill (nearest marker on the same segment) or Rigid Body Fill (for rigid segments) taking advantage of the multiple markers available on each participant. Outliers (*>* 3 SD from the trial mean) were removed per marker and replaced via 1D linear interpolation. Trajectories were smoothed using a 1 s moving-average window. Position magnitude was computed as the Euclidean norm; velocity was derived as the first temporal derivative along the axes separately, with magnitude taken as the Euclidean norm of the resulting vector.

### EEG preprocessing

For both studies, EEG preprocessing followed a validated pipeline implemented in FieldTrip [54] and EEGLAB [55]. Data were bandpass filtered (0.3-95 Hz, Butterworth, third order) and notch filtered (47-53 Hz) to suppress line noise. Channels exhibiting flat signal or exceeding 2.75 SD from the cross-channel amplitude mean were marked as noisy and subsequently interpolated. Artifact Subspace Reconstruction (ASR; [56, 57]; threshold *k* = 10, based on the prior recommendation [58]) removed transient high-variance artifacts. Data were re-referenced to the common average. Ocular artifacts were removed via independent component analysis (ICA) with automatic ICLabel classification [59]. Previously interpolated channels were reconstructed via spherical spline interpolation. Finally, EEG data were downsampled to match their respective behavioural data (study 1: 300 Hz, study 2: 250 Hz).

### Intra- and interpersonal spectral analyses

#### Interpersonal neural synchrony (INS)

Time-aligned EEG from both partners, **X**_1_, **X**_2_ ∈ ℝ^*T*×*C*^, was analysed per channel pair. Complex Morlet wavelet convolution (1-95 Hz, integer frequencies, 7 cycles) yielded analytic signals *W*_1_(*t, c, f*), *W*_2_(*t, c, f*). Instantaneous phase and amplitude were extracted as *ϕ*_1,2_ = ∠*W*_1,2_ and *A*_1,2_ = |*W*_1,2_|.

We computed: Phase-locking value (PLV): 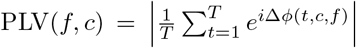, where Δ*ϕ* = *ϕ*_1_ − *ϕ*_2_ and Detrended Amplitude envelope correlation (AmpCorr): Pearson *r*(*A*_1_(*t, c, f*), *A*_2_(*t, c, f*)), Fisher-*z* transformed.

Metrics were averaged across time per dyad and condition (circular mean for circular measures). Cluster-based permutation tests (10,000 permutations) assessed significance against zero (within-condition) and between conditions (paired contrasts; Fisher-*z* for correlations).

#### Contrastive learning of neural-behavioural embeddings

Embeddings were computed for all trials within each dyad using a multi-session CEBRA framework [39], in which data from both participants were concatenated across trials to learn a shared representation of neural-behavioural dynamics.

Two behavioural auxiliary variables were derived per participant per trial block: (1) fingertip velocity (temporal derivative; 0.1-20 Hz bandpass); and (2) submovement velocity (derivative of position; bandpassed at 111111 2-3Hz).

High-dimensional EEG **s**_*t*_ ∈ ℝ^64^ was mapped to a low-dimensional latent space **z**_*t*_ ∈ ℝ^*d*^ (*d* = 3) via a parametric encoder *f*_*θ*_ (3-layer MLP, 32 hidden units). The contrastive objective maximized similarity between neural states linked to similar behavioural contexts (positive pairs: temporal neighbors within Δ*t* = 10 samples) and minimized similarity for dissimilar states (negative pairs: randomly sampled from the dataset). Cosine similarity *ϕ*(**z, z**^*′*^) = **z**^⊤^**z**^*′*^*/*(∥**z**∥**z**^*′*^∥) was employed. The temperature-scaled InfoNCE loss:

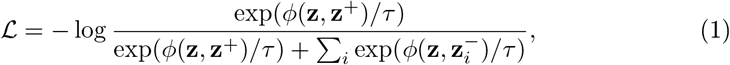

with fixed temperature *τ* = 1. For study 1, training ran for up to 5000 iterations (batch size 512; learning rate 3 × 10^−4^) while study 2 was run for 10000 iterations (to account for the noisy data in study 2 and reach a similar loss between the two studies). Positive and negative pairs were sampled from the full dataset. Model performance was assessed via InfoNCE lower bound on mutual information. Metrics were computed per trial block and dyad.

#### Interpersonal analyses on embeddings

Interpersonal synchrony in embedding space was quantified using two complementary metrics. (1) **Geometric similarity**: Procrustes disparity, corresponding to the squared Procrustes disparity between aligned embedding trajectories. Given **Z**^(1)^, **Z**^(2)^ ∈ ℝ^*T*×*D*^ (*D* = 3), the disparity was defined as 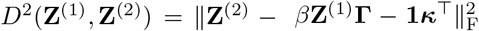, where *β* (scaling), **Γ** (rotation or reflection), and ***κ*** (translation) were optimized to minimize the squared discrepancy. Following standardization, values range from 0 (identical shapes up to the permitted transformations) to 1 (maximal dissimilarity). (2) **Temporal correlation**: Pearson correlations between corresponding embedding dimensions, averaged across dimensions.

### Representational geometry and basis function modeling

To characterize the representational structure of the learned manifolds, we modeled the pairwise Euclidean distance geometry of CEBRA embeddings as a regularized linear combination of nonlinear transformations of the auxiliary kinematic variables. Prior to distance computation, embeddings and kinematic variables were downsampled from 300 Hz to 30 Hz (linear interpolation, factor 10).

Five basis functions were applied to the kinematic variable matrix **X**: Linear (**X**), Absolute (|**X**|), Quadratic (**X**^2^), Cosine (cos **X**), and Sigmoid (*σ*(**X**)). For each basis Φ_*m*_(**X**) and embeddings **Z**, pairwise distance matrices were computed, upper-triangular elements extracted (*k* = 1), and *z*-scored (zero-variance transformations set to zero). The standardized basis distance vectors 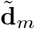 were stacked into predictor matrix **D**_Φ_. Embedding distances **d**_*Z*_ were regressed onto **D**_Φ_ via elastic net (10-fold CV; *L*_1_ ratio grid {0.1, 0.5, 0.7, 0.9, 0.95, 0.99, 1.0}; 5000 maximum iterations). For each L1 ratio, ElasticNetCV automatically generated a path of 100 alpha values spanning *α*_min_*/α*_max_ = 10^−3^, and the combination minimizing the cross-validated error was selected. Cross-validated coefficients *β*_*m*_ were extracted per trial and condition. Dyad-level averaged |*β*_*m*_| were compared via paired *t*-tests with Benjamini-Hochberg FDR correction.

### Dynamics of Latent Spaces

#### Event-locked state-space trajectory extraction

To analyse transient submovement dynamics, continuous state-space trajectories were extracted time-locked to kinematic events. The behavioural state space comprised a 2D manifold of position and submovement velocity; the neural state space comprised the *d* = 3 CEBRA embeddings.

Submovement peaks were detected automatically from velocity profiles as local maxima with prominence *>* 10% of the signal range and minimum separation of 250 ms. The prominence value ensured that any noisy peaks were discarded (especially in study 2 where participants performed spontaneous movements). The minimum separation of 250 ms ensured that no two peaks would have overlapping windows. Non-peak controls were defined as upward or downward signal zero-crossings outside peak-exclusion regions. For each event, a symmetric *±*200 ms window was extracted and resampled to *L* = 200 time points via linear interpolation.

#### Model fitting via autoregressive rolling procedure

Dynamical structure was quantified by fitting a family of low-dimensional attractor models and control models using a rolling autoregressive procedure. Let 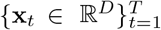 denote a trajectory segment around submovement events, where *D* is the state dimension. Segments were partitioned into *N* trials of fixed length *L*, yielding 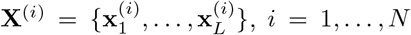. First-order transitions **x**_*t*_ → **x**_*t*+1_ were concatenated across trials to form the dataset *D* = *{*(**x**_*t*_, **x**_*t*+1_)*}*. Model parameters were estimated from *D* via least-squares or moment-based estimators.

##### Point attractor

State updates were modelled as **x**_*t*+1_ = **x**_*t*_ + (**c** − **x**_*t*_)**A**, where **c** was the centroid of the pooled current states and **A** ∈ ℝ^*D*×*D*^ was estimated by least squares from **x**_*t*+1_ − **x**_*t*_ ≈ (**c** − **x**_*t*_)**A**.

##### Trajectory attractor

A template trajectory was defined as 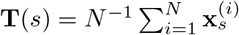. For each state, the nearest template point was identified and the model advanced toward the subsequent template point: **x**_*t*+1_ = **x**_*t*_ + *β*[**T**(*s* + 1) − **x**_*t*_], where *β* was estimated by least squares.

##### Limit cycle

Dynamics were modelled in the first two state-space dimensions. The mean radius and mean circular phase increment were estimated from the pooled transitions, and successive states were generated by advancing the phase by the mean increment at the mean radius.

##### Error-correction control

A scalar correction gain was estimated from transitions toward the global centroid. During rollout, a single fractional correction toward the centroid was applied to the initial state, after which the corrected state was held constant throughout the trajectory.

##### Noise-based control

As a structure-free baseline without explicit temporal dynamics, the model predicted the global centroid at every time step.

##### Model evaluation

Predictions were generated recursively: 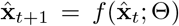 from initial state **x**_1_. This rolling formulation accumulates errors, providing a stringent test of dynamical consistency.

#### Attractor properties: point and trajectory attractors

To characterize geometry and stability, we computed complementary metrics for point and trajectory attractors (Tables 1 and 2). Point-attractor metrics capture global convergence to a fixed equilibrium; trajectory-attractor metrics capture alignment and stability relative to a structured dynamical pathway.

Four dynamical aspects were quantified: (1) **Spatial confinement**: distance to centroid or template; point variance. (2) **Dynamical stability**: local dominant eigenvalue (contraction/expansion). (3) **Metastability**: mean dwell time (persistence near the attractor). (4) **Flow directionality**: flow convergence index (inward/outward flow). For a strong stable point attractor, one expects low distance, low variance, a negative dominant eigenvalue, long dwell time, and high positive flow convergence. These properties were statistically assessed using a two-way repeated-measures ANOVA with peak-type and condition as within-subject factors in study 1, and paired-samples *t*-tests in study 2.

#### Interpersonal coupling metrics

Coordination between partners was quantified on submovement-aligned attractor properties. For trajectory attractors: correlation of template distance between partners; dynamic time warping (DTW) distance. For point attractors: distance between global centroids; distance between endpoints (final state). Metrics were computed only for the winning model from the fitting procedure.

### Granger causality analysis

We applied VAR-based Granger causality [40-42] to quantify directed temporal dependencies between state-space representations. Granger causality measures predictive improvement: *X* Granger-causes *Y* if past values of *X* improve prediction of future *Y* beyond past *Y* alone. Effects are interpreted as directed predictive dependencies, not mechanistic causation [42].

Analyses were performed per dyad, condition, and trial block. Stationarity was verified via Augmented Dickey-Fuller and KPSS tests (all series confirmed stationary). The maximum permitted lag was *p*_max_ = 10 observations (fixed value). Because the input series consisted of ordered event-level features and attractor properties, one lag represented one preceding submovement event rather than a fixed interval in milliseconds. For each variable pair (*X, Y*), we compared:

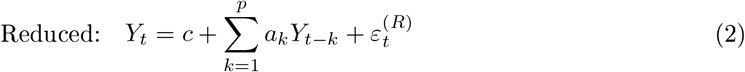

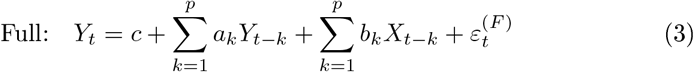

Directed influence: *G*_*X*→*Y*_ = log Var[*ε*^(*R*)^]*/*Var[*ε*^(*F*)^] . Positive values indicate that *X* improves prediction of *Y* . Direction was reversed to estimate *G*_*Y* →*X*_. Lag order *p* was constrained by a maximum lag; models required sufficient observations per lag (at least 2 samples > chosen lag value).

At the dyadic level, we tested bidirectional influences between behavioural and neural state-space trajectories (from CEBRA embeddings) at submovement peaks, assessing whether preceding behavioural-state dynamics predicted subsequent neural population-state dynamics and vice versa.

In complementary single-participant analyses, the same VAR framework was applied to test whether each participant’s behavioural state space predicted subsequent movement parameters (behavioural state as source; movement features as target). This quantified whether behavioural-state dynamics provide predictive information about upcoming movement characteristics above and beyond the movement’s own history. Estimates were computed within trial blocks prior to aggregation.

## Supplementary information

- Supplementary Fig. S1. Global interpersonal measures for study 1.
- Supplementary Fig. S2. Control analyses for the interpersonal alignment of neural embeddings.
- Supplementary Fig. S3. Selection of neural embedding dimensionality.
- Supplementary Fig. S4. Principal component analysis does not reproduce condition-specific interpersonal neural alignment.
- Supplementary Fig. S5. Interpersonal alignment of the kinematic variables used to condition neural embeddings.

## Acknowledgements

This work has received funding from the European Union’s Horizon Europe research and innovation programme under grant agreementNo. 101120727 (PRIMI).

