## Supplementary_information for "Neural and behavioural manifold dynamics align across interacting individuals"

**Contents**


- Supplementary Methods
- Supplementary Fig. 1
- Supplementary Fig. 2
- Supplementary Fig. 3
- Supplementary Fig. 4
- Supplementary Fig. 5

### Supplementary Methods

#### Dyad-level principal component analysis of inter-subject dynamics

For each dyad, multivariate time series from both participants $\mathbf{X}^{(s)} \in \mathbb{R}^{T \times F}$ ($s \in \{1, 2\}$) were jointly normalized: concatenated along the time dimension, feature-wise $z$-scored using pooled $\mu_j$ and $\sigma_j$, then split. A single PCA was fit to the concatenated normalized data, yielding a shared component space. Each participant's data were projected into this space for direct comparison. Inter-subject similarity was quantified via: (1) Procrustes disparity (geometric alignment); and (2) mean Pearson's correlation across corresponding component time series (temporal synchronization). Metrics were computed per block and averaged within conditions.

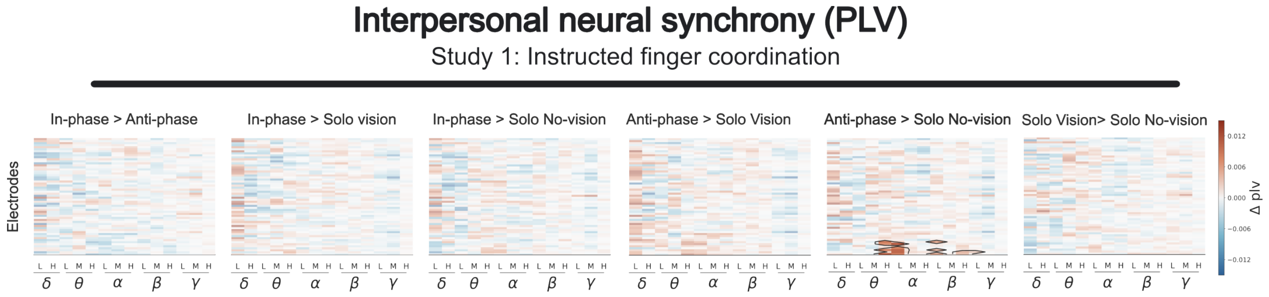


Fig. S1: **Global interpersonal measures for study 1.** Heatmaps of interpersonal neural synchrony (Phase-locking value) for all condition comparisons in study 1. Cluster-based permutation tests revealed significant clusters only for Anti-phase > Solo No-vision condition. No significant effects were found for any other comparison.

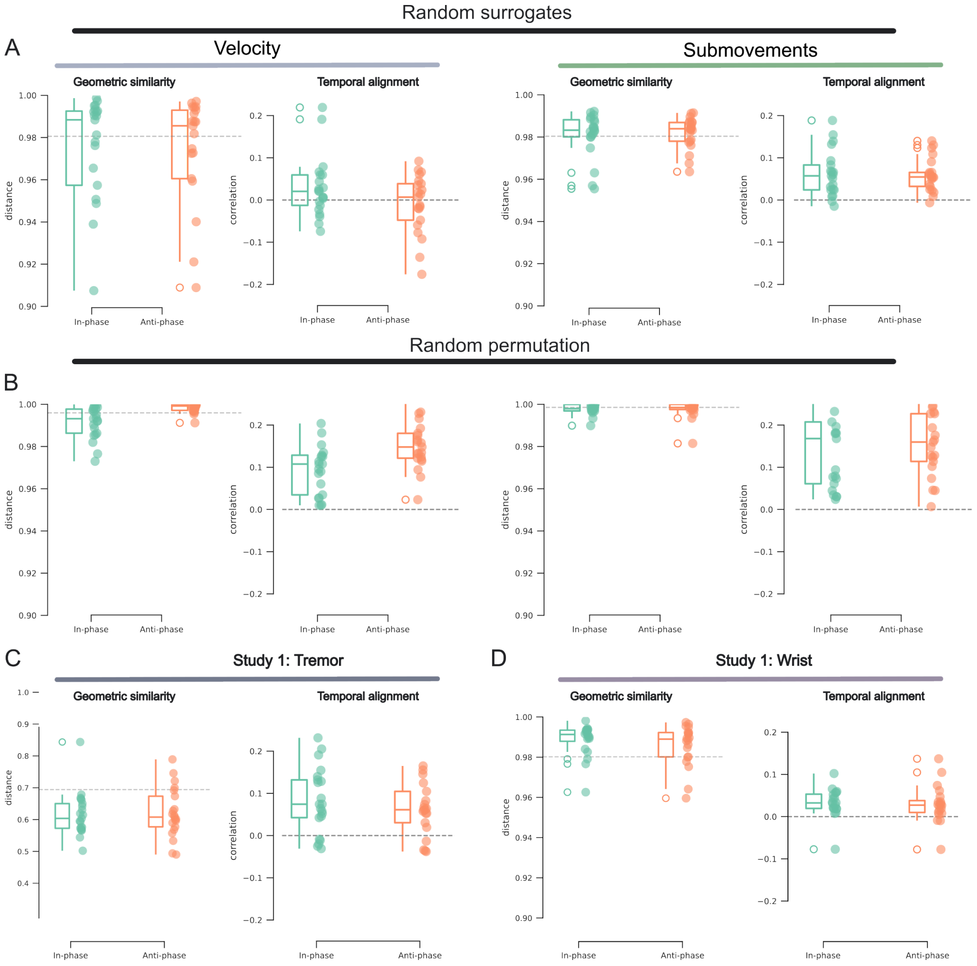


Fig. S2: **Control analyses for the interpersonal alignment of neural embeddings.** **(A)** Trial-surrogate control analysis. Neural embeddings generated in the main analysis were shuffled across interacting dyads, thereby disrupting the correspondence between trials. Box plots show geometric and temporal similarity for the In-phase and Anti-phase conditions at the macro scale (left) and micro scale (right). None of the surrogate analyses reproduced the alignment difference between In-phase and Anti-phase coordination. **(B)** Time-shuffled control analysis for study 1. Neural embeddings were estimated after randomly shuffling the temporal correspondence between the neural and behavioural data. Box plots show geometric and temporal similarity for the In-phase and Anti-phase conditions at the macro scale (left) and micro scale (right). The time-shuffled embeddings did not reproduce the condition-specific interpersonal alignment observed in the original data. **(C)** Frequency-specificity analysis. Embeddings were conditioned on physiological tremor, defined as velocity band-pass filtered at 6–12 Hz, rather than submovement velocity (2–3 Hz). Box plots show geometric and temporal similarity. Physiological tremor did not differentiate the coordination conditions, indicating that the alignment observed in the main analysis was specific to submovements rather than a general property of higher-frequency kinematic fluctuations. **(D)** Effector-specificity analysis. Interpersonal alignment was recomputed using submovement signals derived from the wrist marker. Box plots show geometric and temporal similarity. This analysis did not reproduce the condition-specific effect obtained using the index finger.

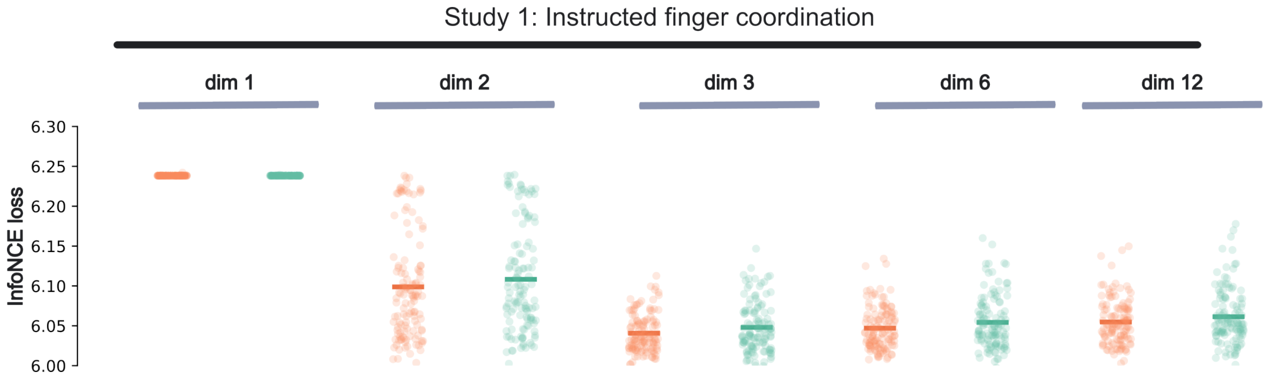


Fig. S3: **Selection of neural embedding dimensionality.** Model performance as a function of embedding dimensionality. CEBRA models were trained using latent spaces of increasing dimensionality, and performance was evaluated using the InfoNCE loss (lower values correspond to better fits). A three-dimensional embedding provided the most parsimonious representation; dimensionalities greater than three yielded equivalent condition effects. Dots represent individual dyads; horizontal lines indicate the group mean.

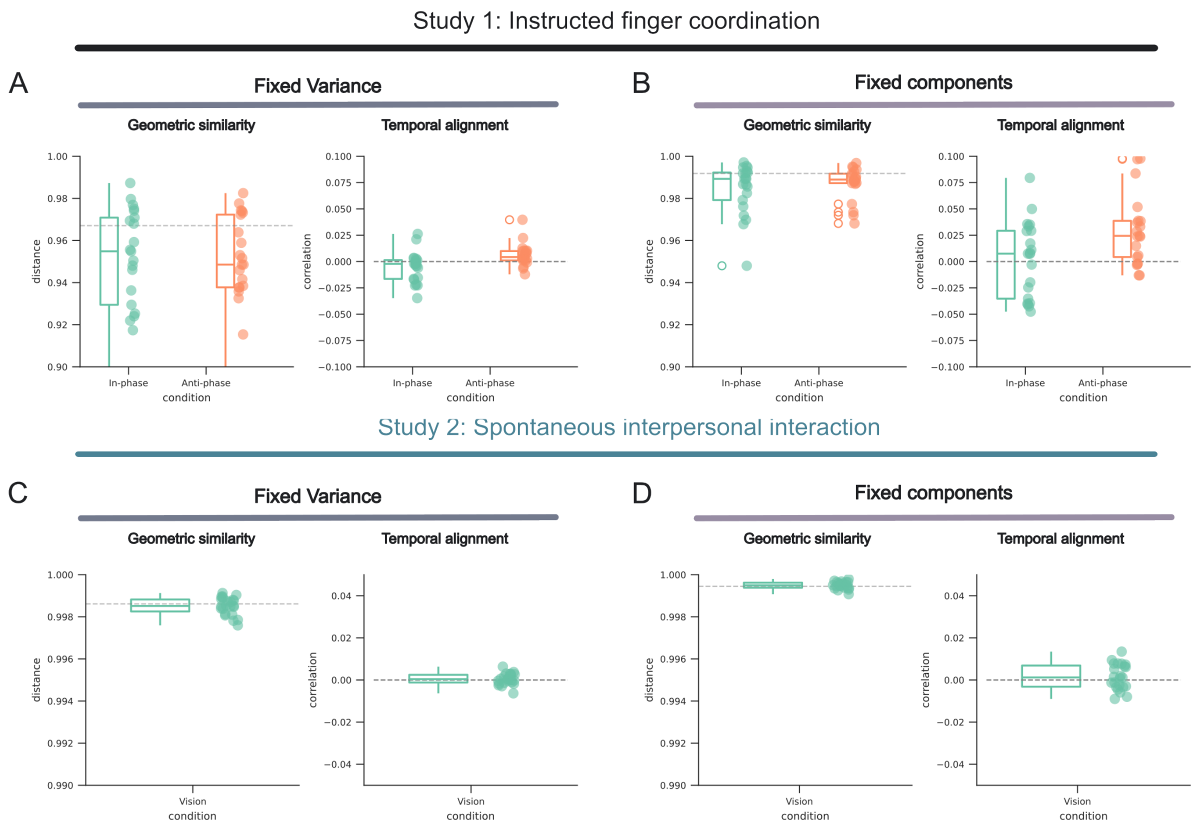


Fig. S4: **Principal component analysis does not reproduce condition-specific interpersonal neural alignment.** **(A)** Study 1 results obtained using principal component analysis (PCA), retaining the number of components required to explain 90% of the variance. Geometric similarity between partners' PCA trajectories was quantified using Procrustes disparity (left), and temporal alignment was quantified using the Pearson's correlation between corresponding principal-component time series (right). **(B)** Study 1 results obtained using PCA with the number of retained components fixed at three. Geometric similarity between partners' PCA trajectories was quantified using Procrustes disparity (left), and temporal alignment was quantified using the Pearson's correlation between corresponding principal-component time series (right). **(C)** Study 2 results obtained using PCA, retaining the number of components required to explain 90% of the variance. Geometric similarity between partners' PCA trajectories was quantified using Procrustes disparity (left), and temporal alignment was quantified using the Pearson's correlation between corresponding principal-component time series (right). **(D)** Study 2 results obtained using PCA with the number of retained components fixed at three. Geometric similarity between partners' PCA trajectories was quantified using Procrustes disparity (left), and temporal alignment was quantified using the Pearson's correlation between corresponding principal-component time series (right). Unlike the behaviourally conditioned CEBRA embeddings, the unconstrained linear PCA embeddings did not reliably distinguish In-phase from Anti-phase coordination in study 1 or Vision from No-vision interaction in study 2.

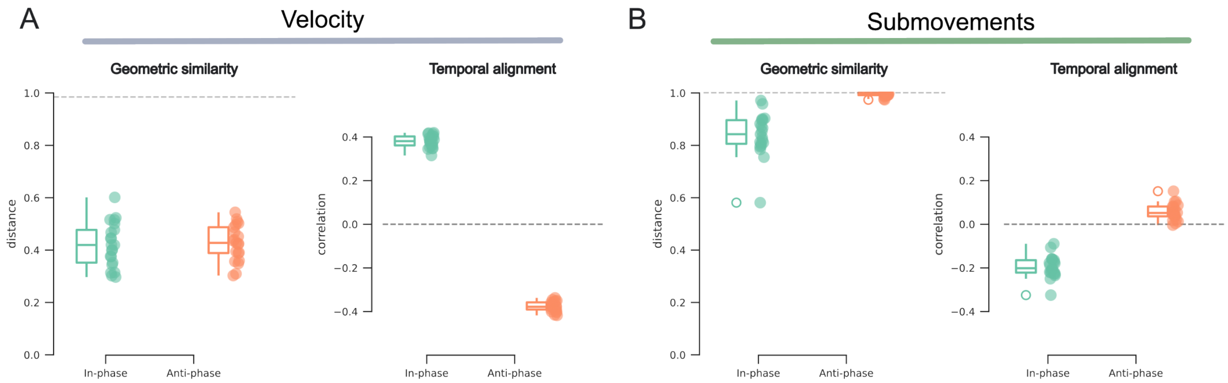


Fig. S5: **Interpersonal alignment of the kinematic variables used to condition neural embeddings. (A)** Macro-scale. Geometric similarity (left) and temporal similarity (right) of study 1 kinematic trajectories, quantified using Procrustes disparity and Pearson's correlations respectively for velocity. In-phase movements showed positive temporal alignment at the macroscopic velocity scale, whereas Anti-phase movements showed negative temporal alignment. **(B)** Micro-scale. Geometric similarity (left) and temporal similarity (right) of study 1 kinematic trajectories, quantified using Procrustes disparity and Pearson's correlations respectively for submovement velocity. At the submovement scale, the temporal relationship followed the corresponding micro-scale coordination structure. These behavioural results reproduce the condition-dependent geometry and temporal organization observed in the behaviourally conditioned neural manifolds.
